# Two distinct lipid transporters together regulate invasive filamentous growth in the human fungal pathogen *Candida albicans*

**DOI:** 10.1101/2022.08.30.505893

**Authors:** Miguel A. Basante-Bedoya, Stéphanie Bogliolo, Rocio Garcia-Rodas, Oscar Zaragoza, Robert A. Arkowitz, Martine Bassilana

## Abstract

Flippases transport lipids across the membrane bilayer to generate and maintain asymmetry. The human fungal pathogen *Candida albicans* has 5 flippases, including Drs2, which is critical for filamentous growth and phosphatidylserine (PS) distribution. Furthermore, a *drs2* deletion mutant is hypersensitive to the antifungal drug fluconazole and copper ions. We show here that such a flippase mutant also has an altered distribution of phosphatidylinositol 4-phosphate [PI(4)P], and ergosterol. Analyses of additional lipid transporters, *i.e.* the flippases Dnf1-3, and all the oxysterol binding protein (Osh) family lipid transfer proteins, *i.e.* Osh2-4 and Osh7, indicate that they are not critical for filamentous growth. However, deletion of Osh4 alone, which exchanges PI(4)P for sterol, in a *drs2* mutant can bypass the requirement for this flippase in invasive filamentous growth. In addition, deletion of the lipid phosphatase Sac1, which dephosphorylates PI(4)P, in a *drs2* mutant results in a synthetic growth defect, suggesting that Drs2 and Sac1 function in parallel pathways. Together, our results indicate that a balance between the activities of two different classes of lipid transporters regulates invasive filamentous growth, *via* PI(4)P. In contrast, deletion of *OSH4* in *drs2* does not restore growth on fluconazole, nor on papuamide A, a toxin that binds PS in the outer leaflet of the plasma membrane, suggesting that Drs2 has additional role(s) in plasma membrane organization, independent of Osh4. As we show that *C. albicans* Drs2 localizes to different structures, including the Spitzenkörper, we sought to determine if a specific localization of Drs2 is critical for different functions, using a synthetic physical interaction approach to restrict/stabilize Drs2 at the Spitzenkörper. Our results suggest that Drs2 plasma membrane localization is critical for *C. albicans* growth on fluconazole and papuamide A, but not for invasive filamentous growth.

## Introduction

Polarized growth is an essential process that is regulated, in particular, by cooperative interactions between key establishment proteins and specific lipids at the plasma membrane (PM). For instance in the baker’s yeast *Saccharomyces cerevisiae*, dynamic nanoclusters of the Rho-GTPase Cdc42 are regulated by multivalent interactions between its sole activator Cdc24, the scaffold protein Bem1 and anionic lipids, including PS [1–3]. Similarly, in the fission yeast *Schizosaccharomyces pombe*, the localization and function of the two essential Rho-GTPases, Cdc42 and Rho1, depend on polarized PS distribution [4]. The importance of anionic lipids, and PS in particular, for cellular function and signaling encompasses kingdoms from fungi, to mammals [5] to plants [6]. In addition, PS has been reported to be critical for virulence in a broad spectrum of microbial pathogens (reviewed in [7, 8]). In particular, in the human fungal pathogen *Candida albicans*, the PS synthase Cho1, which is conserved in fungi but does not have an ortholog in Human, is required for virulence in a murine candidiasis model [9]. A *cho1* deletion mutant also exhibits increased exposure of β(1,3)-glucan *via* up-regulation of cell wall MAPK cascades, facilitating its detection by innate immune cells [10]. PS is synthesized in the endoplasmic reticulum (ER) and its cellular distribution is regulated both by lipid transfer proteins (LTPs), which function at contact sites between the ER and the target cellular compartments, and lipid transporters, such as flippases that establish PS asymmetry between membrane leaflets (reviewed in [11]).

Lipid flippases are P4-ATPases, only found in eukaryotes, where they are similar in domain structures from fungi to Humans (reviewed in [12]). These proteins actively transport phospholipids from the external/luminal to cytoplasmic membrane leaflets and play an important role in polarized growth (reviewed in [13]). There are 14 P4-ATPases in Human and only 5 in the yeast *S. cerevisiae* (Dnf1-3, Drs2, Neo1), with most of them regulated by non-catalytic subunits from the Cdc50/Lem3 family. For example, in *S. cerevisiae*, Cdc50 forms a functional complex with Drs2 [14, 15] and very recently, the cryo-electron microscopy structure of different conformations of this complex was resolved at 2.8 to 3.7 Å [16]. This complex, which is well characterized, both *in vitro* and *in vivo,* primarily transports PS [17, 18]. The role of lipid flippases on membrane curvature and in trafficking has been extensively studied in *S. cerevisiae*, as well as in mammals (reviewed in [19]). In *S. cerevisiae*, Neo1, which is the sole essential flippase [20], localizes to the Golgi and endomembranes, Drs2 and Dnf3 to the trans-Golgi network (TGN) and Dnf1-2 to the PM [17, 21, 22]. A recent study indicates that Dnf3 can also be found at the PM in a cell-cyle dependent fashion, where it regulates, together with Dnf1-2, *S. cerevisiae* pseudohyphal growth [23]. Furthermore, in response to pheromone, it was shown that Dnf1, Dnf2 and Dnf3 localize to the schmoo tip, while Drs2 remains at the Golgi [24]. Interestingly, in the filamentous fungi *Aspergillus nidulans* and *Fusarium graminearum,* it was shown that the Drs2 homolog, DnfB, localizes primarily to a cluster of vesicles at the hyphal apex, called Spitzenkörper (SPK), which is characteristic of fungi that grow in a filamentous form [25, 26]. Together, these data indicate that flippases can localize to different distinct compartments in specific conditions of polarized growth.

LTPs and more specifically oxysterol-binding protein (OSBP)-related proteins (ORPs), are also important for membrane lipid composition, *via* non-vesicular traffic (reviewed in [27]), including for PS distribution (reviewed in [11]). LTPs can bind specific ligands such as PI(4)P, PS and sterol. For example, in *S. cerevisiae*, there are 7 oxysterol-binding homology (Osh) proteins, among which Osh6/Osh7 transports PS from cortical ER (cER) and PM [28], in counter-exchange with PI(4)P [29]. On the other hand, another Osh protein, Kes1, which is homolog to Osh4 and transports sterol in counter-exchange with PI(4)P [30], was proposed to act antagonistically with Drs2 to regulate the sterol distribution between PM and internal membranes in *S. cerevisiae* [31].

Filamentous fungi are highly polarized organisms, in which the role of flippases has been investigated. In *A. nidulans*, the homologs of Dnf1 and Drs2, *i.e.* DnfA and DnfB, regulate growth and PS asymmetry [25], while in *Magnaporthe grisea,* Mg*APT2*, one of the 4 aminophospholipid translocase (APT) encoding genes related to Drs2, is required for plant infection but not filamentous growth [32]. In *F. graminearum*, flippases play redundant as well as distinct roles in vegetative growth, where FgDnfA is critical, stress response, reproduction and virulence [26, 33, 34]. In the human fungal pathogen *C. albicans*, deletion of either Dnf1 or Dnf2 results in moderate increase sensitivity to copper [35]. The *drs2* deletion mutant is hypersensitive to copper [35], as well as to the antifungal drug fluconazole [36]. Deletion of Drs2 also results in altered PS distribution and impaired filamentous growth [36], and that of its Cdc50 subunit in altered filamentous growth and reduced virulence in a murine candidiasis model [37]. Here, we sought to determine more specifically the role of Drs2 in morphogenesis and whether PM PS asymmetry is critical for this process.

Our results show that in response to serum, of the four flippases Dnf1-3 and Drs2, only Drs2 is critical for filamentous growth, with Dnf2 having a minor role in this process. Filamentous growth was largely recovered in the *drs2* mutant upon deletion of the LTP Osh4, but not by that of other Osh proteins (Osh2, Osh3 and Osh7). Furthermore, our results demonstrate that the distribution of PS and of the phosphatidylinositol phosphate PI(4)P, which are both altered in the *drs2* mutant, is restored upon deletion of *OSH4*. These data indicate that the requirement for flippase activity across the lipid bilayer during filamentous growth can be specifically bypassed by lipid exchange between membrane compartments.

## Results

### Drs2 has a unique role in *C. albicans*

In *A. nidulans*, the Drs2 homolog DnfB, is not critical for hyphal growth [25]. In contrast, in *C. albicans*, Drs2 is required for filamentous growth, whereas budding growth is largely unaffected (mean doubling time of 90 min, compared to 80 min in the wild-type strain) [36]. We investigated the importance of other flippases, *i.e.* Dnf1, Dnf2 and Dnf3, using loss of function mutants that we generated (Supplementary Figure S1A). As illustrated in Figure 1A, a *drs2* deletion mutant was not invasive on serum-containing media, and reintroduction of a copy of *DRS2* in this mutant restored invasive growth (Figure 2A). In contrast, deletion of *DNF1* or *DNF3* did not alter serum-induced invasive growth, while deletion of *DNF2* resulted in reduced invasive growth, which was further reduced in a double *dnf1dnf2* deletion mutant. Over-expression of *DNF2* restored invasive growth in the *dnf2* and *dnf1 dnf2* mutants, but not in the *drs2* mutant, suggesting that Dnf2 and Drs2 do not functionally overlap in this invasive growth process. Upon serum-induced hyphal growth in liquid media (Figures 1B-D), the *dnf1, dnf2* and *dnf3* deletion mutants produced hyphae similar to the wild-type cells, with hyphal formation in the *dnf1 dnf2* cells somewhat reduced. The average hyphal filament length was, nonetheless, slightly reduced in *dnf2* cells as well as in *dnf1 dnf2* (14 ± 3 μm and 13 ± 3 μm, respectively) compared to the WT, *dnf1* and *dnf3* cells (20 ± 4 μm, 18 ± 4 μm and 20 ± 4 μm, respectively). These results are in agreement with very recent data, which also show that both a *dnf2* and a *dnf1 dnf2* mutant exhibit reduced invasive filamentous growth in response to the nutrient poor spider media, another inducer of filamentous growth [38]. These data indicate that, in contrast to the *drs2* deletion mutant, *dnf1*, *dnf2* and *dnf3* deletion mutants undergo filamentous growth similar to the wild-type cells, albeit with a reduced efficiency for *dnf2*. Furthermore, Figure 1E shows that the *dnf1-3* mutants also grew similar to the wild-type cells in the presence of the cell wall perturbants calcofluor white (CFW) and congo red (CR), and the antifungal drugs, caspofungin (Caspo) and fluconazole (FCZ). In contrast, the *drs2* mutant was specifically hypersensitive to CFW and FCZ [36]. The growth defect of *drs2* cells on FCZ could result from a mislocalization of multi-drug transporters, such as Cdr1 [39]. Figure 1F shows that, although a Cdr1-GFP fusion protein was detected at the PM in *drs2* cells, 79 ± 8% of the cells had internal Cdr1 signal, in contrast to wild-type cells, suggesting that PM targeting of MDR is partially impaired in *drs2* cells. Together, these data indicate that Drs2 has a unique role in *C. albicans*.

**Figure 1:**
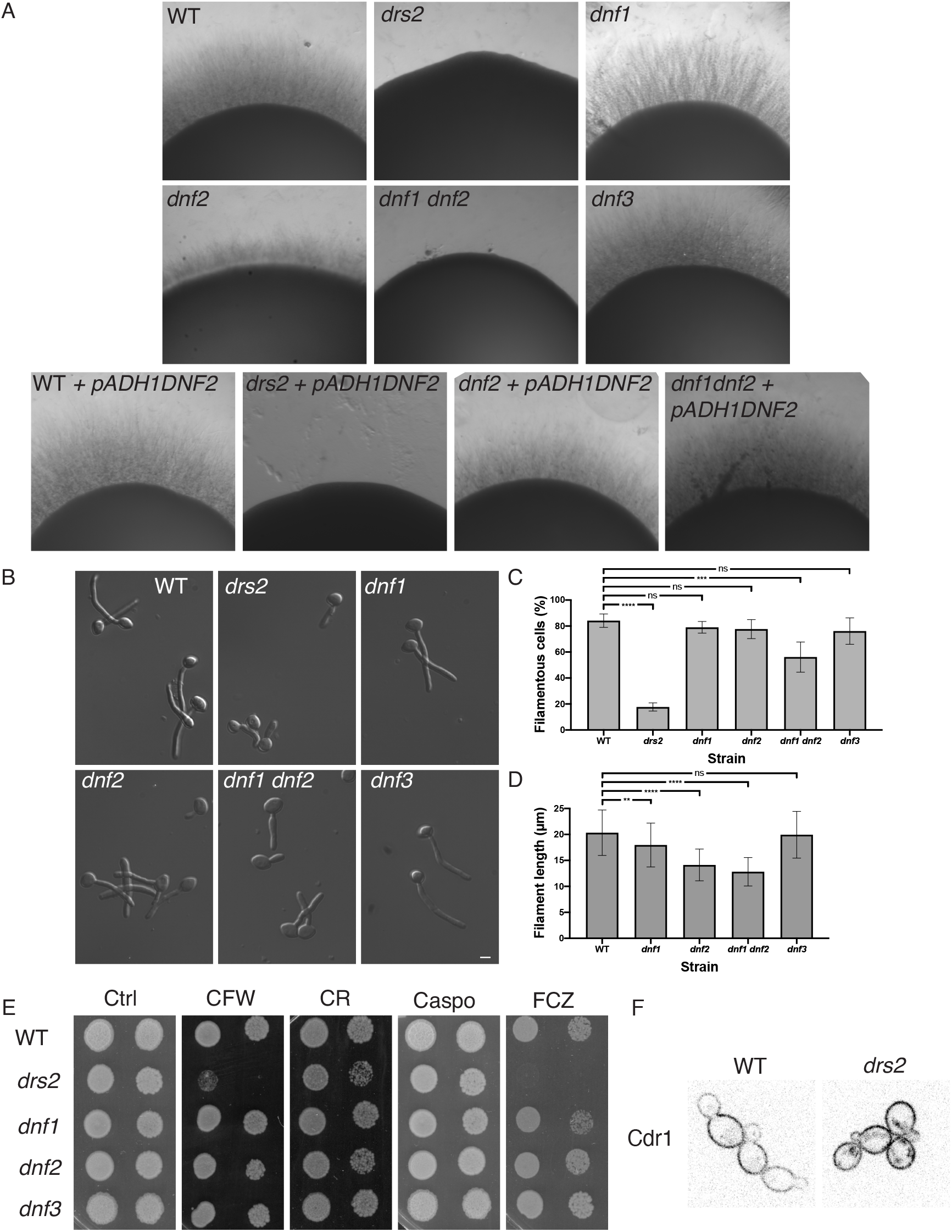
Drs2 has a unique role in *C. albicans.* A) Invasive filamentous growth cannot be restored by overexpression of *DNF2* in the *drs2/drs2* mutant. The indicated strains, WT (PY4861), *drs2/drs2* (*drs2*, PY3375), *dnf1/dnf1* (*dnf1*, PY6235), *dnf2/dnf2* (*dnf2*, PY5814), *dnf3/dnf3* (*dnf3*, PY5801), *dnf1/dnf1 dnf2/dnf2* (*dnf1 dnf2*, PY6400), WT + *pADH1DNF2* (PY5005), *drs2/drs2* + *pADH1DNF2* (*drs2* + *pADH1DNF2,* PY5003)*, dnf2/dnf2* + *pADH1DNF2* (*dnf2 + pADH1DNF2*, PY5919), and *dnf1/dnf1 dnf2/dnf2* + *pADH1DNF2* (*dnf1 dnf2* + *pADH1DNF2*, PY5922), were grown on agar-containing YEPD with serum and images were taken after 6 days. Similar results were observed in 2 independent experiments. B) *DRS2* is specifically required for hyphal growth in response to serum. Cells from the indicated strains were incubated with serum for 90 min at 37°C. Bars are 5 µm. C) and D). Graphs represent the percentage of hyphae (C) and the filament length (D) in the indicated strains grown as in 1B. The percentage of hyphae was 84 ± 5%, 79 ± 4%, 78 ± 7%, 76 ± 10%, 56 ± 12% and 18 ± 3% for wild-type and the *dnf1, dnf2*, *dnf3, dnf1 dnf2* and *drs2* deletion mutants, respectively (average of 3 experiments with *n* ∼ 150 cells each). The filament length was measured from the junction between cell body and filament (error bars indicate the mean +/- the SD of 3 experiments, *n* ∼ 50 cells each). **, *P* < 0.01; ***, *P* < 0.0005; **** *P* < 0.0001; ns, not significant. (E). The *drs2* mutant has specific increased susceptibility to fluconazole and calcofluor white. Serial dilutions of indicated strains were spotted on YEPD media (Ctrl) containing 25 μg/ml calcofluor white (CFW), 400 μg/ml Congo red (CR), 125 ng/ml caspofungin (Caspo) or 5 μg/ml fluconazole (FCZ). Images were taken after 2 days at 30°C. Similar results were observed in 2 independent experiments. (F). PM targeting of the multidrug ABC transporter Cdr1 is altered in *drs2* cells. Central z-sections of representative WT and *drs2/drs2* cells expressing Cdr1-GFP are shown.

**Figure 2:**
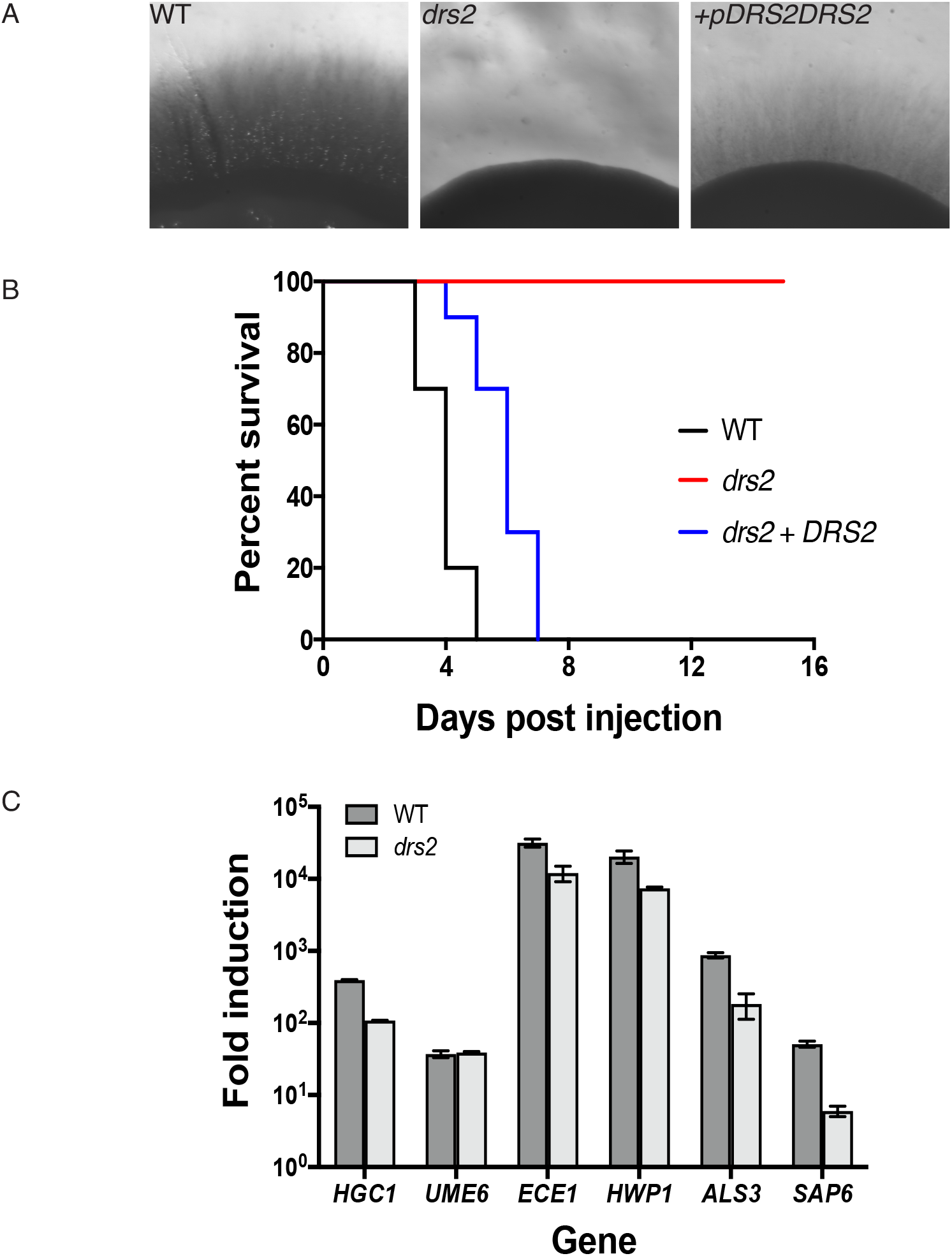
Drs2 is critical for virulence in a mouse model of systemic candidiasis. A) Reintroduction of *DRS2* complements the invasive growth defect of the *drs2* mutant. The indicated strains WT (PY4861), *drs2/drs2* (*drs2*, PY3375), and *drs2/drs2 + pDRS2DRS2* (*drs2 + DRS2*, PY5042), were grown as in Figure 1A. B) The *drs2* mutant has attenuated virulence in a mouse model of hematogenously disseminated candidiasis. Balb/C mice (*n* = 10) were injected with an inoculum (5 x 10^5^ cells) of the indicated strains and the survival was assessed. C) Hyphal specific gene are induced in *drs2* cells upon serum exposure. mRNA and cDNA were prepared from the indicated strains grown 120 min in the presence of serum. *HGC1*, *ECE1*, *HWP1*, *ALS3* and *SAP6* transcripts were determined by qRT-PCR using primer pairs described in [89], and *UME6* transcripts were determined using UME6.pTm/UME6.mTm (89 bp) primer pair. Bars indicate the mean ± SD (*n* = 3 determinations). Similar results were observed in an additional biological replicate. NAD-linked Glyceraldehyde-3-phosphate dehydrogenase (*TDH3*) transcript levels were used for normalization.

### Drs2 is critical for hyphal extension after septin ring formation

In response to serum, *drs2* cells appear to initiate filamentous growth, although they were unable to form hyphae [36]. We examined this mutant in a murine model for systemic candidiasis and Figure 2B shows that the *drs2* mutant is nonvirulent, compared to the wild-type and complemented strains. At the transcriptional level, hyphal growth is controlled by the hyphal specific cyclin *HGC1*, which is further regulated by the transcription factor *UME6* [40]. Both *hgc1* and *ume6* deletion mutants are defective in hyphal extension and attenuated for virulence in a mouse infection model [41, 42]. Quantitative RT-PCR analyses in Figure 2C show that, upon serum exposure, *HGC1* and *UME6* were up-regulated in the *drs2* mutant compared to budding cells (>100-fold for *HGC1* and 40-fold for *UME6*), with a level of induction of *HGC1* slightly reduced compared to that of the wild-type cells. Similarly, *ECE1*, which encodes the peptide toxin candidalysin and whose expression correlates with cell elongation [43], *HWP1*, which encodes a hyphal cell wall protein associated with hyphal development [44] and *ALS3,* which encodes an agglutinin-like (Als) adhesin [45], were all up-regulated in the *drs2* mutant upon serum exposure, with levels slightly reduced (3- to 5-fold) compared to the control cells. Only the induction of *SAP6,* which encodes a secreted acid protease [46], was substantially reduced (∼ 10-fold) in *drs2*. Together, these data indicate that the *drs2* mutant hyphal growth defect is unlikely due to the modest reduction of *HGC1* induction.

To further characterize the *drs2* mutant at the molecular level, we used time-lapse microscopy to follow cells expressing fluorescent reporters for different cellular compartments. Figure 3A illustrates representative time courses of hyphal growth both in WT and *drs2* cells expressing a fluorescently tagged septin Cdc10 [47]. Measurements of the diameters of filament compartments, apical (pre-septum) and distal (post-septum) to the septin ring (Figure 3B), show that, while it remained constant in wild-type cells (1.8 ± 0.2 µm), the diameter increased in the apical compartments of *drs2* cells by about 40% (to 2.5 ± 0.3 µm). This increase in diameter was associated with a reduced extension rate (on average 0.09 µm/min in *drs2* cells, compared to 0.34 µm/min in WT cells; Figure 3C). These data are consistent with a defect in polarized growth in the *drs2* mutant after the first septin ring forms. In agreement with this, we observed that, upon filament extension, active Cdc42, visualized with the CRIB (Cdc42 Rac1 Interactive Binding domain) reporter [48], became depolarized in *drs2* cells and the SPK (visualized with the myosin light chain Mlc1, [49, 50]), was not maintained at the filament tip following cell division, compared to the control cells (Figures 3D and 3E). Furthermore, comparison by time-lapse microscopy of the wild-type and *drs2* cells expressing both reporters for active Cdc42 and the Spitzenkörper, indicates that depolarization of Cdc42 occurred prior to the SPK delocalization from the filament tip. Together, these data indicate that, subsequent to septin ring formation, the *drs2* mutant is unable to redirect or reinitiate growth to the apex, which ultimately results in growth arrest and/or pseudohyphal growth.

**Figure 3:**
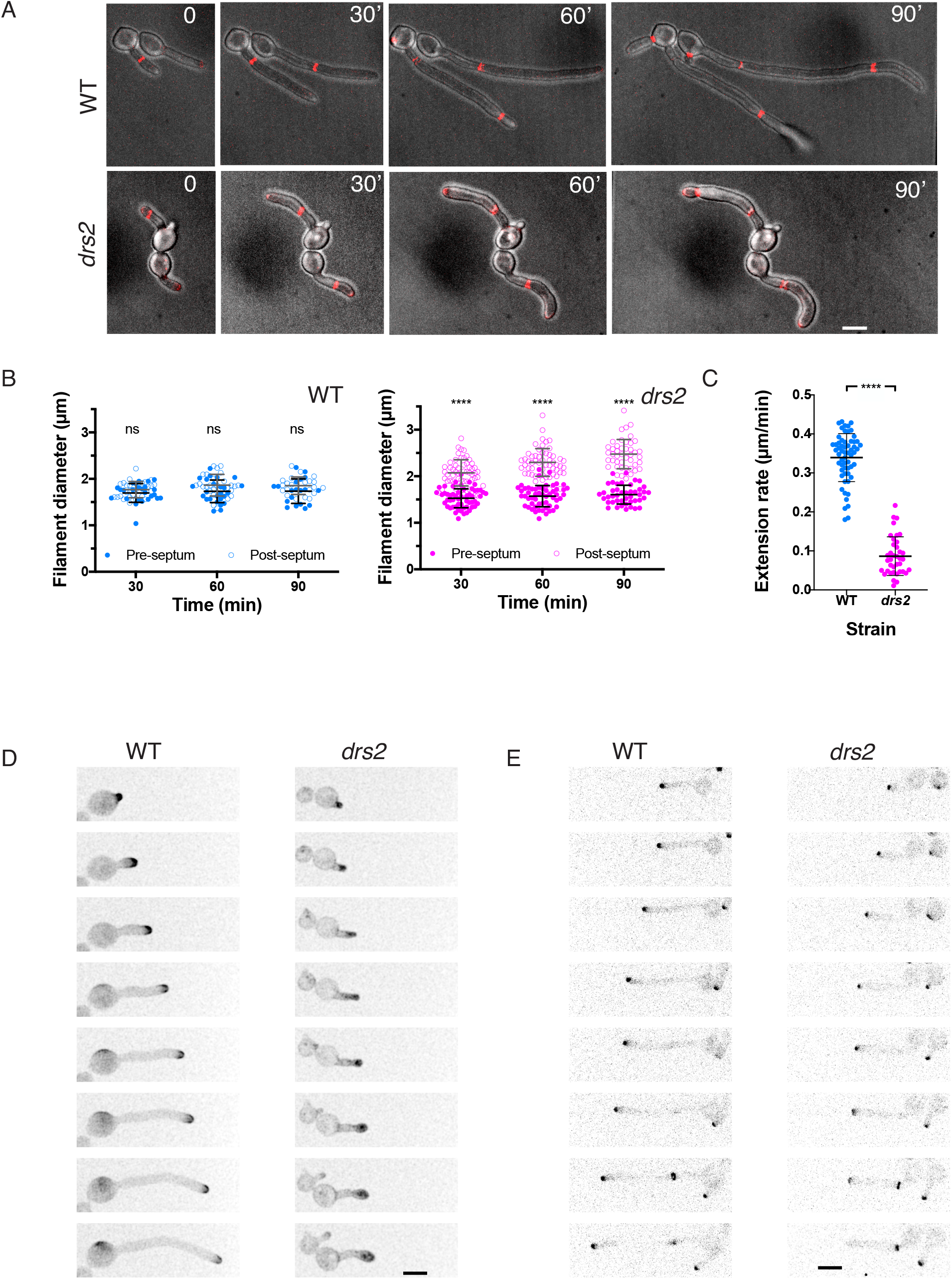
Drs2 is critical for maintaining polarized filament extension. A) Both cell morphology and filament extension rate are altered in the *drs2* mutant, after septin ring formation. Time lapse of wild-type (WT, PY5613) and *drs2/drs2* (*drs2*, PY5615) cells expressing Cdc10-mScarlet, incubated in the presence of serum. Images were taken every 10 min and merges between DIC and sum projections of 21 z-sections are shown. Bars are 5 µm. B) Graphs represent the filament diameter before (open circles) and after (solid circles) the first septin ring, as a function of the times from the first image in which the septin ring is observed in wild-type (blue) and *drs2* (magenta) cells. Means ± SD of 25-50 cells are shown. **** *P* < 0.0001; ns, not significant. C) The graph shows the filament extension rate in wild-type (blue) and *drs2* (magenta) cells. Means ± SD of 30-60 cells are shown. D) and E) Polarized growth is altered in the *drs2* mutant. D) Time lapse of wild-type (WT, PY2263) and *drs2* cells (*drs2*, PY4972) expressing CRIB-GFP, incubated in the presence of serum. Images were taken every 10 min and maximum projections of 21 z-sections are shown. E) Time lapse of wild-type (WT, PY5349) and *drs2/drs2* cells (*drs2*, PY5218) expressing Mlc1-mScarlet, incubated in the presence of serum. Images were taken every 5 min and sum projections of 23 z-sections are shown.

### Drs2 localizes to the hyphal tip

In *S, cerevisiae*, Drs2 localizes to the late Golgi during budding [20] and mating [24], while its homolog in *A. nidulans*, DnfB, localizes within the SPK core [25]. To examine the distribution of Drs2 in *C. albicans*, we generated a strain that expresses a *DRS2-GFP* fusion, which was functional (Figure 4A). Given that the homolog of Dnf1-2 in *A. nidulans,* DnfA, also localizes to the Spitzenkörper, albeit to the outer layer macrovesicles compared to DnfB [25], we also generated a strain that expressed a functional *DNF2-GFP* fusion for comparison (Figure 4A). Figure 4B shows that Drs2 was restricted to the apical region of the hyphal tip, as well as in internal structures, likely Golgi cisternae by analogy with *S. cerevisiae* [20]. Dnf2 was essentially localized at the apical cortex, similar to its localization in *S. cerevisiae* at bud tips and mating projections [24]. Both Dnf2 and Drs2 co-localized with a PM marker (a prenylated RFP fusion, RFP-CtRac1, [51]), although Drs2 localized to a more restricted region of the hyphal tip than Dnf2 (Figure 4C). Drs2 also partially co-localized with the SPK marker, Mlc1 (Figure 4D), which could be due to the proximity (less than 100 nm) of this structure to the PM [52]. Together, these data indicate that Drs2 and Dnf2 have distinct distributions at the filament apex.

**Figure 4:**
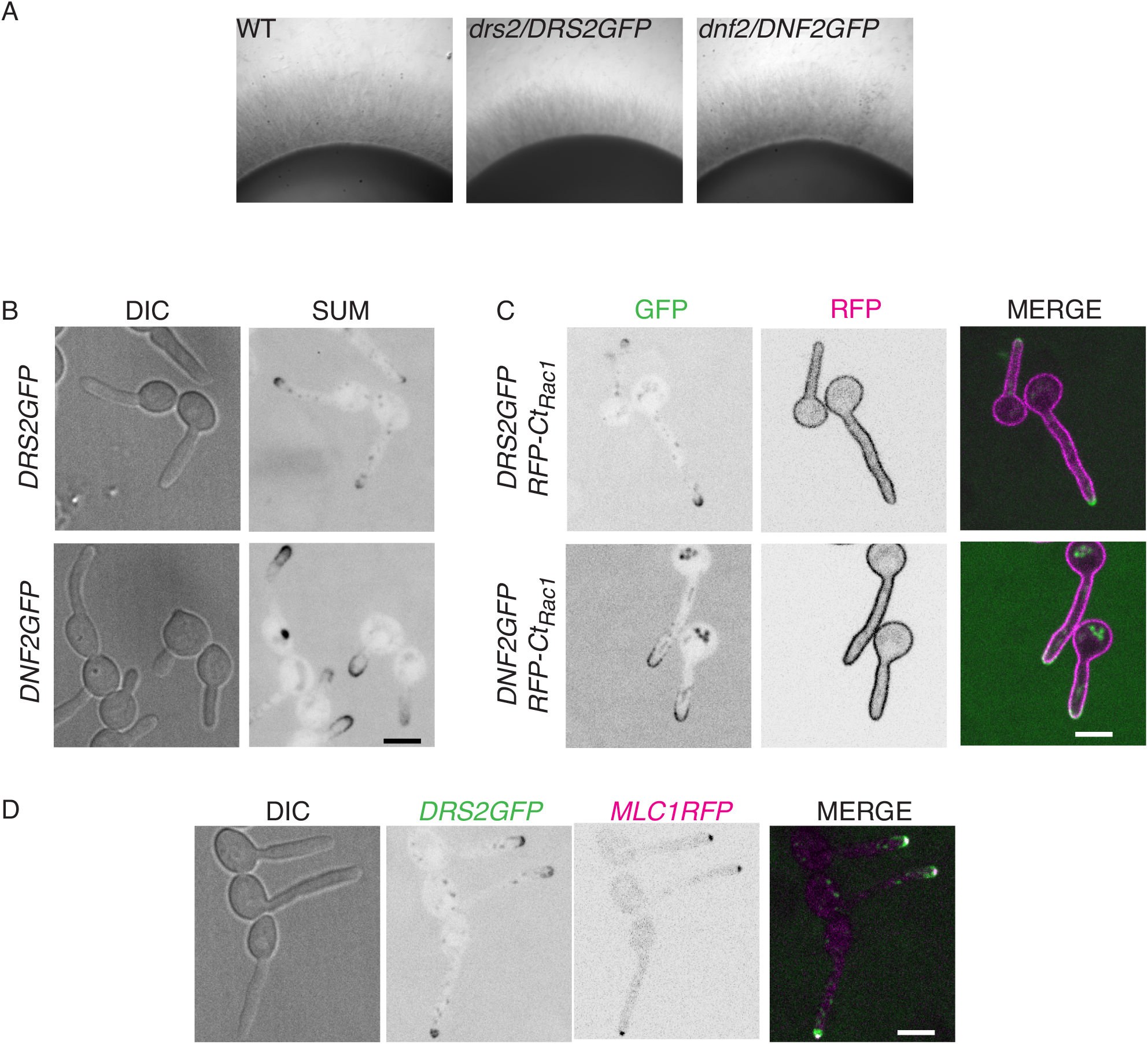
Drs2 localizes preferentially to the filament apex. A) The Drs2-GFP and Dnf2-GFP fusions are functional. The indicated strains, WT (PY4861), *drs2*/*DRS2GFP* (PY4665) and *dnf2/DNF2GFP* (PY5746) were grown as in Figure 1A. B) and C) Drs2 and Dnf2 localize differently to the filament apex. Sum projections of 16 z-sections of representative cells expressing Drs2-GFP and Dnf2-GFP after 90 min serum induction are shown (B). Central z-sections and merge of representative cells expressing mScarlet-CtRac1, together with either Drs2-GFP (PY6241) or Dnf2-GFP (PY6239), after 90 min serum induction are shown (C). D) Drs2 partially overlaps with Mlc1. Central z-section and merge of representative cells expressing Drs2-GFP together with Mlc1-mScarlet (PY4928), after 90 min serum induction are shown. Bars are 5 µm.

### The *drs2* mutant is altered for PI(4)P distribution

Drs2 is a P4-ATPase that flips PS selectively across the lipid bilayer *in vitro* and *in vivo* in *S. cerevisiae* [17, 18]. In *C. albicans*, using the reporter Lactadherin C2 (LactC2) [53–55], we observed that the distribution of PS was altered during hyphal growth in the *drs2* mutant, as the reporter was visible as intracellular punctae [36]. To determine the impact of the *DRS2* deletion on the distribution of other lipids, shown to be critical for hyphal growth, such as the phosphatidylinositol phosphates PI(4)P and PI(4,5)P_2_ [56, 57], as well as ergosterol [58], we used specific fluorescent reporters. The distribution of PI(4,5)P_2_ appears to be similar in wild-type cells and *drs2* filamentous cells (Figure 5A), yet the distribution of PI(4)P was substantially less polarized in the mutant, compared to the wild-type or complemented strains (Figure 5B). This depolarization is further illustrated by the graph in Figure 5B, which shows the relative concentration of PI(4)P, as a function of filament length. In contrast, PI(4)P at the Golgi was largely unaffected in *drs2*, compared to WT cells, both during filamentous growth (Figure 5C), and budding growth (mean of 7.1 ± 2.8 Golgi cisternae per cell in *drs2* cells compared to 7.8 ± 2.4 in the wild-type control, Figure 5D). Together, these data indicate that deletion of *DRS2* results in altered distribution of PM PI(4)P, but not PI(4,5)P_2_.

**Figure 5:**
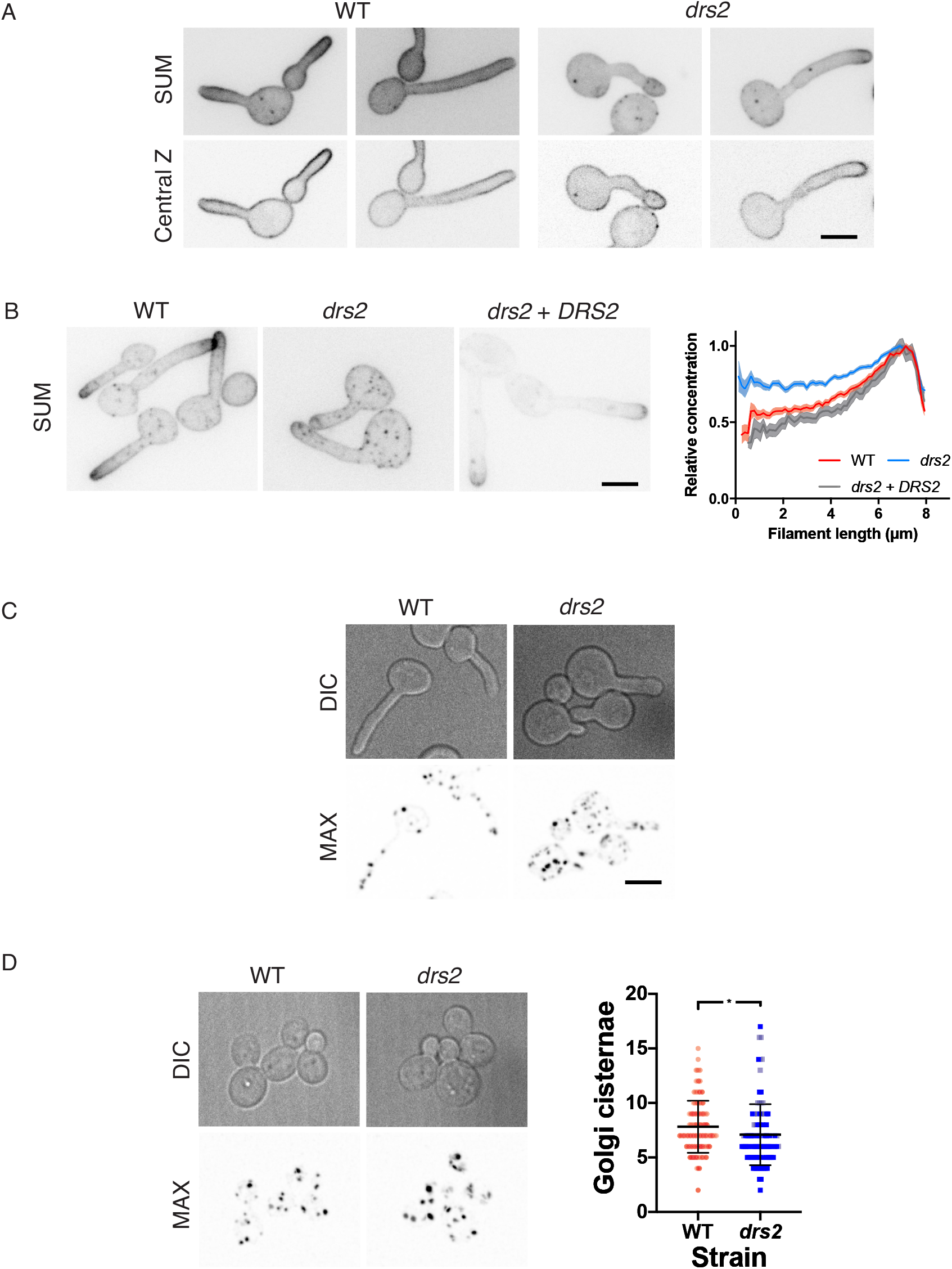
Plasma membrane PI(4)P distribution is altered in the *drs2* mutant. A) PM PI(4,5)P_2_ distribution is not altered in the *drs2* mutant. Wild-type (PY1206) and *drs2/drs2* cells (*drs2*, PY4050) expressing GFP-(PH^PLCδ1^)_2_-GFP were incubated for 60 or 90 min, respectively, in the presence of serum. Sum projections (21 z-sections) and central z-sections of representative cells are shown. B) PM PI(4)P distribution is altered in the *drs2* mutant. The indicated cells, WT (PY5619), *drs2/drs2* (*drs2*, PY5568) and *drs2/drs2* + *pDRS2DRS2* (*drs2 + DRS2*, PY6407) expressing GFP-(PH^OSH2[H340R]^)_2_-GFP were incubated for 90 min in the presence of FCS. Sum projections of (21 z-sections) of representative cells are shown. The graph illustrates the means ± the SEM of the relative concentration of PM PI(4)P as a function of filament length, normalized to the maximal signal for each cell (*n* = 25-60 cells). C) and D) The number of Golgi cisternae is not substantially affected in the *drs2* mutant. DIC and maximum projections (21 deconvolved z-sections) of representative WT (PY2578) and *drs2/drs2* cells (*drs2*, PY3873) expressing FAPP1^[E50A,H54A]-^GFP incubated in the presence of serum are shown (C). The number of Golgi cisternae per cell was determined in budding cells of the indicated strains from maximum projections of deconvolved images (21 z-sections). (D). Bars indicate the mean ± the SD of 3 independent biological samples (*n* = 100 cells and ∼700-800 cisternae for each strain). * *P* < 0.05.

Using filipin staining, it was shown that membrane sterols are highly concentrated at the apex during *C. albicans* hyphal growth, with such a polarization not observed in budding and pseudohyphal cells [58]. We examined sterol distribution, using both filipin staining and the genetically encoded biosensor D4H [59, 60]. Figure 6A shows that sterols are highly concentrated at the apex of WT hyphal cells, irrespective of the reporter used. In contrast, in the *drs2* mutant, the D4H reporter preferentially labeled internal structures and reintroduction of a copy of *DRS2* restored the ergosterol distribution to the PM (Figure 6B). Together, these data indicate that deletion of *DRS2* results not only in altered distribution of PM PS, but also of PI(4)P and ergosterol.

**Figure 6:**
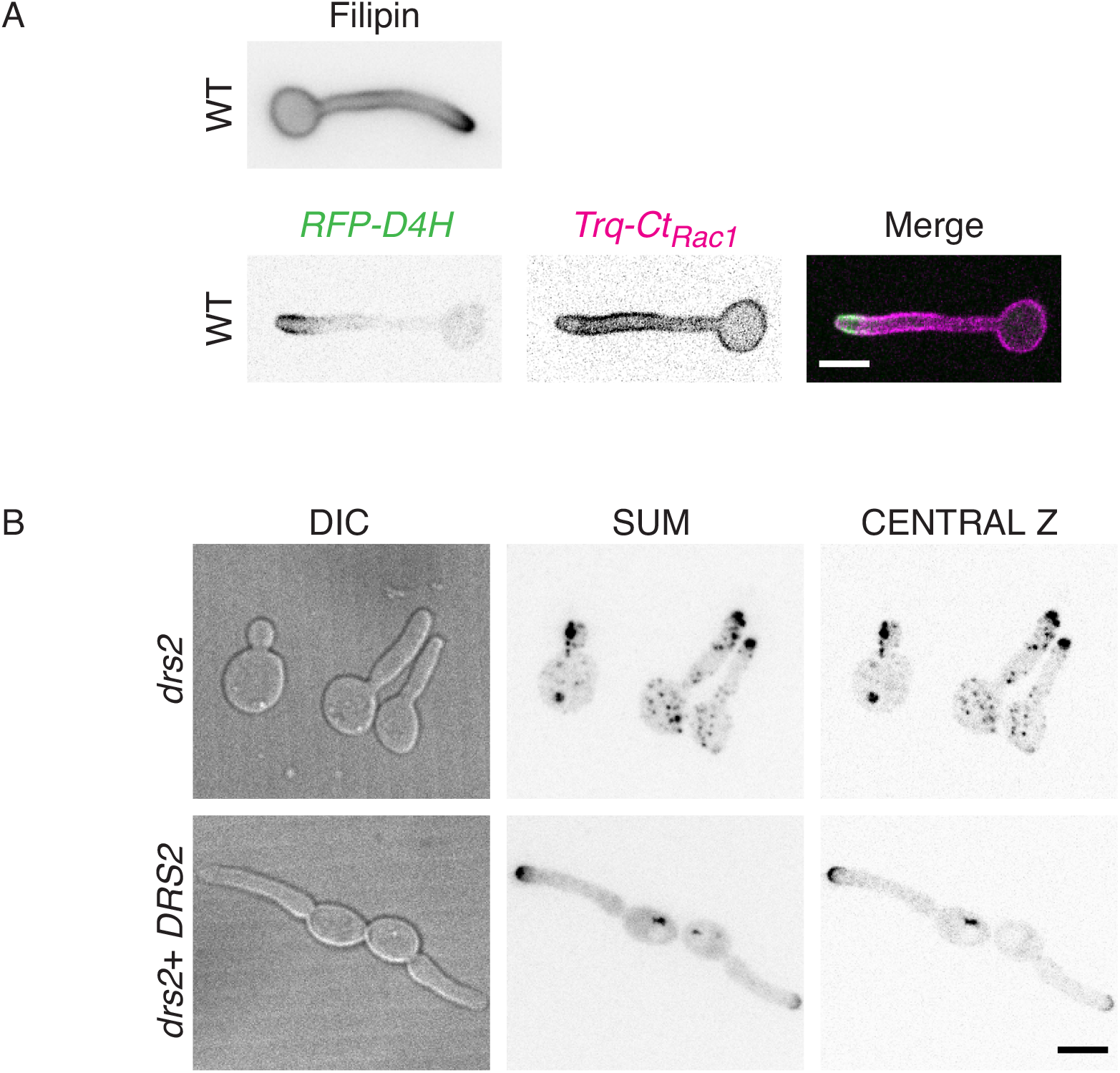
Distribution of ergosterol is altered in the *drs2* mutant. A) The ergosterol reporters Filipin and D4H localize similarly at the apex of the filament in wild-type cells. Top panel: Wild-type cells (PY4861) were induced serum prior to staining with filipin, as described [58]. Images were taken with a wide-field fluorescence microscope. Bottom panel: Wild-type cells expressing mScarlet-D4H together with Trq-CtRac1 (PY6237) were induced with serum as in Fig. 1B and images were taken with a spinning disk confocal microscope and central z-section, as well as merge images, are shown. B) Ergosterol distribution is altered in the *drs2* mutant. Images of *drs2/drs2* cells (*drs2*, PY6083) and *drs2/drs2* + *pDRS2DRS2* cells (*drs2 + DRS2*, PY6218) expressing mScarlet-D4H were taken as in Figure 6A bottom panel.

### Deletion of *OSH4* in the *drs2* mutant recovers invasive filamentous growth and plasma membrane PI(4)P distribution

LTPs can bind specific ligands such as PI(4)P, PS and sterol and, in *S. cerevisiae*, it was shown that Osh6/Osh7 transports PS in counter-exchange with PI(4)P [29], while Kes1 transports sterol in counter-exchange with PI(4)P [30]. *C. albicans* has 4 Osh proteins (Osh2-4 and Osh7), with Osh4 and Osh7 sharing ∼ 60% identity with their *S. cerevisiae* counterparts, however their function is unknown in this organism. Note that the genes encoding *OSH7* (called *OBPA* and *OBPALPHA*) are located at the mating-type-like locus (MTL), similar to those encoding the Golgi phosphatidylinositol kinase Pik1 (*PIKA* and *PIKALPHA*).

To investigate the importance of Osh proteins in *C. albicans* invasive hyphal growth, we first generated *osh4* and *osh7* deletion mutants, as well as *drs2 osh4 and drs2 osh7* double deletion mutants (Supplementary Figure S1B), and examined these mutants for invasive hyphal growth in response to serum. While deletion of *OSH4* or *OSH7* alone did not alter invasive growth, deletion of *OSH4,* but not *OSH7,* in the *drs2* mutant recovered invasive growth to a level similar to that of WT cells (Figure 7A). In serum-containing liquid media, we also observed a recovery of hyphal growth upon deletion of *OSH4* in the *drs2* mutant (Figure 7B). In this *drs2 osh4* mutant, hyphal length was only slightly reduced compared to that of the wild-type and *osh4* cells (17 ± 3 µm, compared to 19 ± 1 µm for WT and *osh4* cells). To confirm the specificity of the *OSH4* deletion in recovering invasive growth in the *drs2* mutant, we also generated *drs2 osh2* and *drs2 osh3* double deletion mutants (Supplementary Figure S2A). *C. albicans OSH3* has been reported to be important for invasive growth in nutrient poor media (Spider media), but not in the presence of serum [61]. Similarly, we observed that mutants deleted for either *OSH2* or *OSH3* alone grew invasively in the presence of serum (Supplementary Figure S2B). In contrast to *OSH4* deletion, deletion of either *OSH2* or *OSH3* did not recover the *drs2* invasive growth defect, although colonies from the *drs2 osh3* mutant were somewhat crenelated compared to the control strains (Supplementary Figure S2B). Furthermore, while the *osh2* and *osh3* mutants formed hyphae similar to that of the wild-type (83 ± 5%, 73 ± 9% and 86 ± 6% for the WT, *osh2* and *osh3* strains, respectively), the percent of hyphae in the *drs2 osh2* and *drs2 osh3* mutants was similar to that of *drs2* (8 ± 6% and 21 ± 7%, respectively, compared to 19 ± 8% in *drs2*) (Supplementary Figure S2C). These data indicate that while Osh proteins (Osh2-4 and Osh7) are not required for serum-induced invasive filamentous growth, deletion of O*SH4* specifically bypasses the *drs2* requirement in this process.

**Figure 7:**
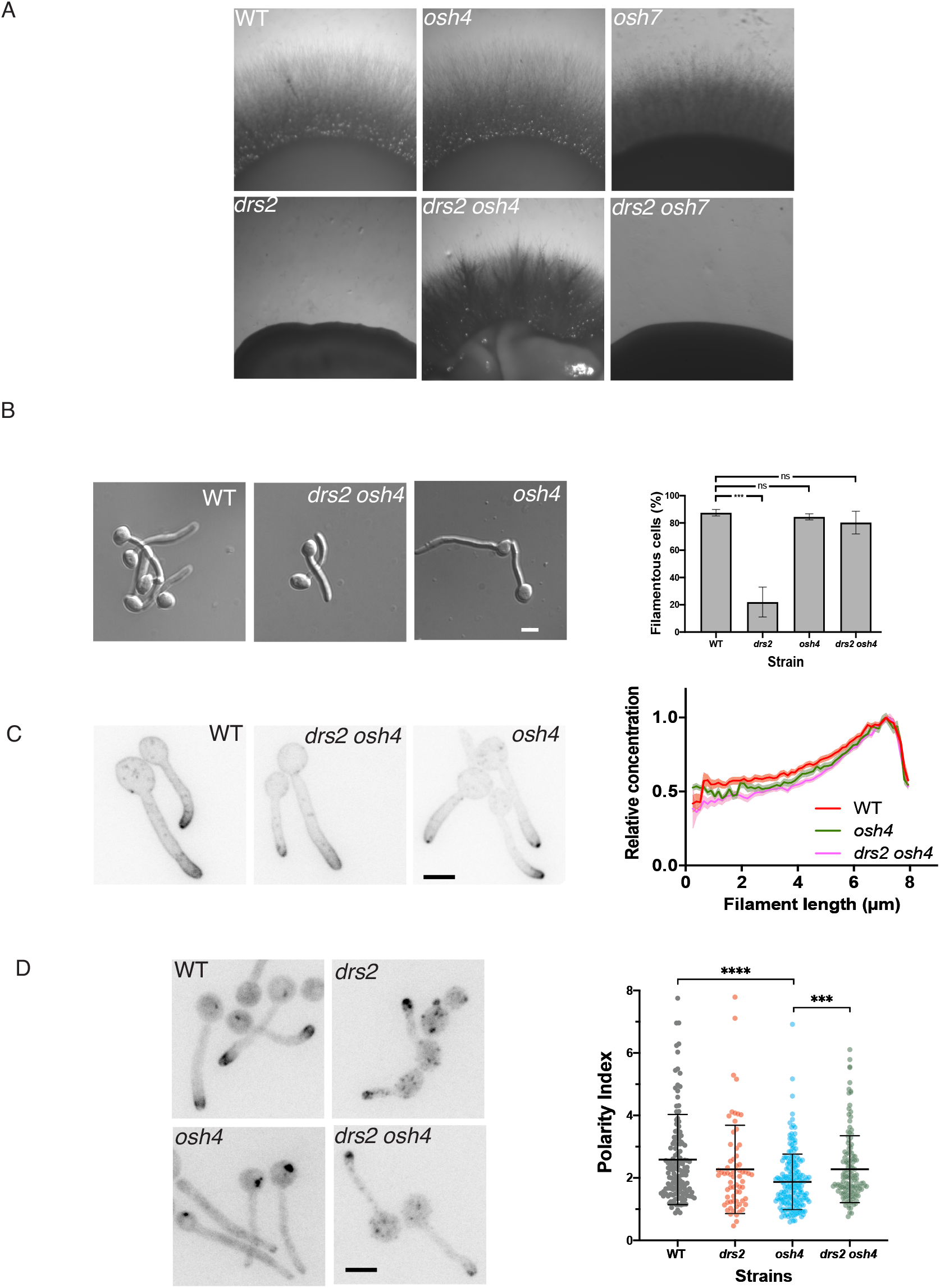
Deletion of *OSH4* rescues hyphal invasive growth and PI(4)P distribution in the *drs2* mutant. A) Invasive filamentous growth is specifically restored in the *drs2* mutant upon deletion of *OSH4*. The indicated strains, WT (PY4861), *osh4/osh4* (*osh4*, PY3974), *osh7/osh7* (*osh7*, PY5256), *drs2/drs2* (*drs2*, PY3375), *drs2/drs2 osh4/osh4* (*drs2 osh4*, PY5539) and *drs2/drs2 osh7/osh7* (*drs2 osh7*, PY5297), were grown as in Figure 1A and images were taken after 6 days. Similar results were observed in 2 independent experiments. B) Hyphal growth is restored in the *drs2* mutant upon deletion of *OSH4.* Cells from the indicated strains were incubated with serum as in Fig. 1B. Bars are 5 µm. The graph represents the percentage of hyphae in the indicated strains, calculated as in Figure 1C. The percentage of hyphae was 82 ± 8%, 80 ± 6% and 76 ± 9% for the wild-type, *osh4* and *drs2 osh4* cells, respectively. ***, *P* < 0.0005; ns, not significant. C) PI(4)P distribution is restored in the *drs2* mutant upon *OSH4* deletion. Indicated cells expressing GFP-(PH^OSH2[H340R]^)_2_-GFP, WT (PY5619), *drs2/drs2 osh4/osh4* (*drs2 osh4*, PY5630) and *osh4/osh4* (*osh4*, PY5626) were incubated as in Figure 5B, with sum projections of representative cells shown. The graph illustrates the means ± the SEM of the relative concentration of PM PI(4)P as a function of filament length, normalized to the maximal signal for each cell (*n* = 100 cells). D) *OSH4* deletion does not substantially restore ergosterol distribution in the *drs2* mutant. Images of the indicated strains expressing mScarlet-D4H, WT (PY6037), *drs2/drs2 osh4/osh4* (*drs2 osh4*, PY6286) and *osh4/osh4* (*osh4*, PY6076) were taken as in Figure 6C. The graph represents the ergosterol tip polarization, *i.e.* the ratio of apical to sub-apical D4H signals, in the indicated strains.

To gain further insight into the mechanisms underlying the recovery of hyphal growth in the *drs2 osh4* mutant, we examined whether deletion of *OSH4* also restored the lipid distribution in the *drs2* mutant. Figure 7C shows that the distribution of PI(4)P was polarized in the *drs2 osh4* double mutant, similar to that in wild-type and *osh4* cells. Similarly, deletion of *OSH4* in the *drs2* mutant significantly restored PS distribution, as internal punctae were absent in *drs2 osh4* cells, similar to the wild-type, complemented and *osh4* strains (Supplementary Figure S3A). Quantification of the LactC2 signal in the *drs2* mutant shows that the fraction of PS was reduced at the PM and increased internally, compared to wild-type and *osh4* cells, resulting in a decreased ratio of PM to intracellular signals (Supplementary Figure S3B). Deletion of *OSH4* in *drs2* significantly increased this ratio, with an increase in PM signal, as well as a decrease in internal signal (Supplementary Figure S3B). In contrast, the overall distribution of ergosterol appears similar in *drs2* and *drs2 osh4*, with the presence of a number of internal punctae, compared to wild-type and *osh4* cells (Figure 7D). The ergosterol tip polarization, determined as the apical versus sub-apical D4H signal, was also similar in these two mutants and not substantially different from that of wild-type cells. Notably, the ergosterol tip polarization was significantly reduced in the *osh4* mutant alone, which exhibits hyphal growth similar to wild-type cells, suggesting that there is not a direct correlation between ergosterol tip polarization and hyphal growth. Together, these data indicate that Drs2 *per se* is not required for hyphal invasive growth, but rather that a balance in the activities of Drs2 and Osh4 regulate this process, likely *via* regulation of the PM PI(4)P gradient.

### Deletion of both *DRS2* and *SAC1* results in a synthetic growth defect

To further examine the importance of the PM PI(4)P gradient, we generated mutants deleted for the lipid phosphatase Sac1, which dephosphorylates PI(4)P, and is responsible for regulating PM PI(4)P, both in yeast and mammalian cells [62, 63]. In *C. albicans,* Sac1 is critical for the steep PM PI(4)P gradient [56] and hyphal growth maintenance [56, 64]. In *S. cerevisiae*, deletion of either *KES1* or *SAC1* is synthetically lethal in cells largely devoid of ER-PM contact sites [65], and cold-sensitive growth of a *drs2* deletion mutant is partially suppressed by deletion of either *SAC1* or *KES1* [31]. As deletion of *OSH4* restored PI(4)P polarized distribution in *drs2*, as well as hyphal growth, we investigated the effect of *SAC1* deletion in the *drs2* mutant and RT-PCR confirmed the absence of *DRS2* and/or *SAC1* in the respective mutants (Figure 8A). Figures 8B and 8C show that budding growth was altered in such a *drs2 sac1* mutant, with a 3-fold increased doubling time, compared to the WT, *drs2* and *sac1* strains. Furthermore, in serum-containing liquid media, while the *sac1* mutant formed filamentous cells, albeit shorter than the wild-type cells (Figure 8D; [56, 64]), the *drs2 sac1* mutant was unable to form filamentous cells, even after 270 min incubation. Together, these data show that *DRS2* and *SAC1* genetically interact in *C. albicans*, indicating that they function in parallel pathways.

**Figure 8:**
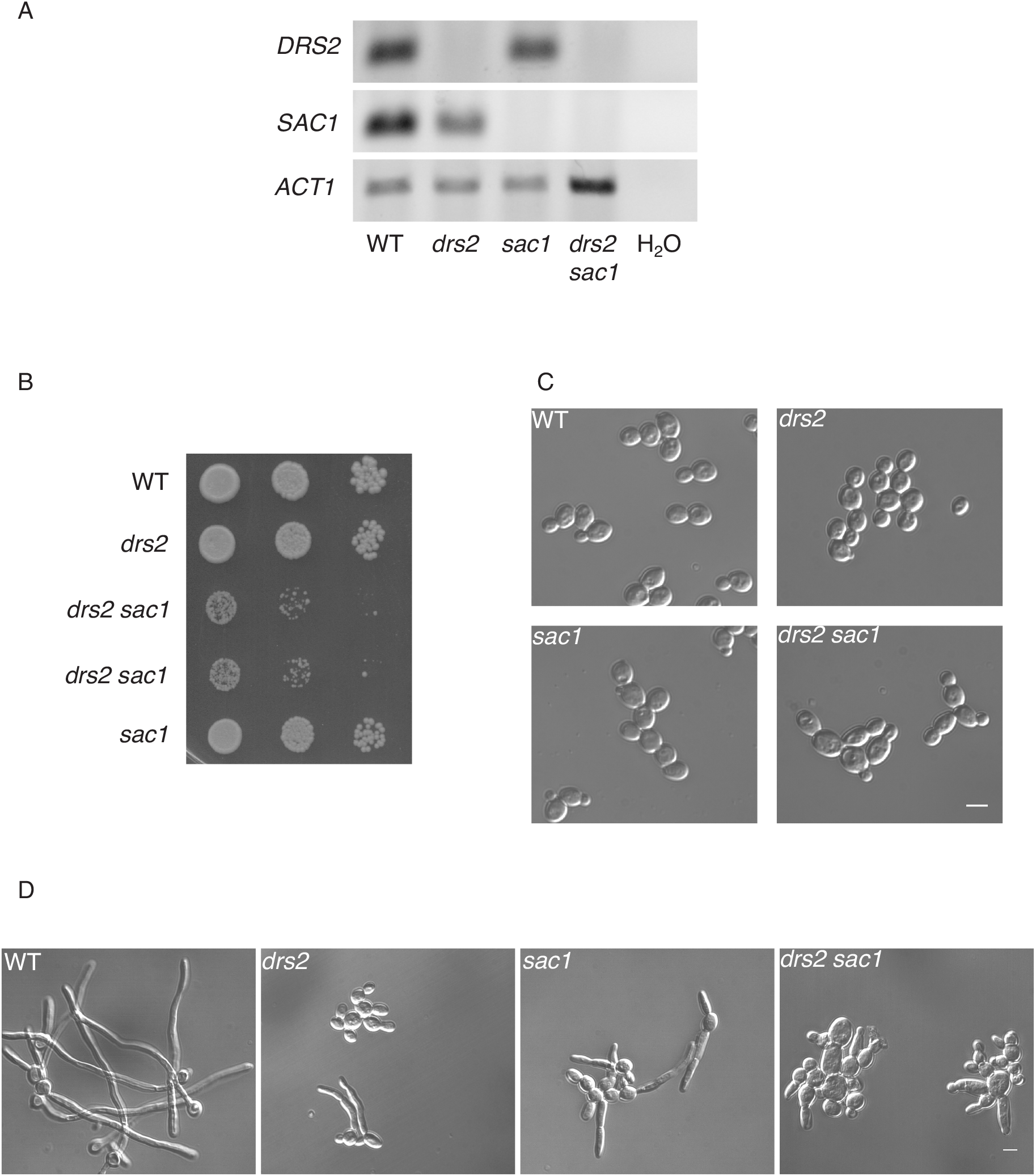
*DRS2* and *SAC1* genetically interact during *C. albicans* development. A) *DRS2* and *SAC1* transcript levels. Transcripts of the indicated strains, as in Fig. 9B, were determined by RT-PCR, using *SAC1*.pTm/*SAC1*.mTm (90 bp) and *DRS2*.pTm/*DRS2*.mTm (62 bp) primer pairs. B) Deletion of both *DRS2* and *SAC1* results in a synthetic growth defect. Serial dilutions of indicated strains, WT (PY4861), *drs2/drs2* (*drs2*, PY3375), *drs2/drs2 sac1/sac1* (*drs2 sac1*, PY6431 & PY6432) and *sac1/sac1* (*sac1*, PY6436), were spotted on YEPD and images were taken after 3 days. Similar results were observed in 2 independent experiments. C) Morphology of *drs2 sac1* budding cells. Cells from the indicated strains WT (PY4861), *drs2/drs2* (*drs2*, PY3375), *drs2/drs2 sac1/sac1* (*drs2 sac1*, PY6432) and *sac1/sac1* (*sac1*, PY6436), were grown to exponential phase in YEPD. The doubling time was 270 min for *drs2 sac1*, compared to 90 min for WT. D) The *drs2 sac1* mutant does not generate hyphae in response to serum. Indicated strains, as in Fig. 9B, were grown in the presence of serum for 180 min. At 90 min, the percent of filamentous cells was 83 ± 5%, 19 ± 8%, 38 ± 4% and 2 ± 1% for the WT, *drs2*, *sac1* and *drs2 sac1* strains, respectively; *n* ∼ 100 cells. Bars are 5 µm.

### Drs2 is important for plasma membrane organization

In addition to a defect in filamentous hyphal growth, the *drs2* mutant also exhibits a hypersensitivity to CFW and FCZ, compared to wild-type cells (Figure 1E). We investigated whether a balance in the activities of Drs2 and Osh4 is also critical for growth on these compounds, and, while deletion of *OSH4* in the *drs2* mutant recovered the growth defect on CFW, it did not recover that on FCZ (Figure 9A). Furthermore, as the *drs2* mutant was reported to be hypersensitive to copper ions [35], we investigated whether deletion of *OSH4* could restore growth in this condition. Figure 9B shows that both *drs2* and *drs2 osh4* mutants were similarly reduced for growth on CuSO4, compared to wild-type and *osh4* cells, indicating that Drs2 has specific roles that are not linked to Osh4 activity.

**Figure 9:**
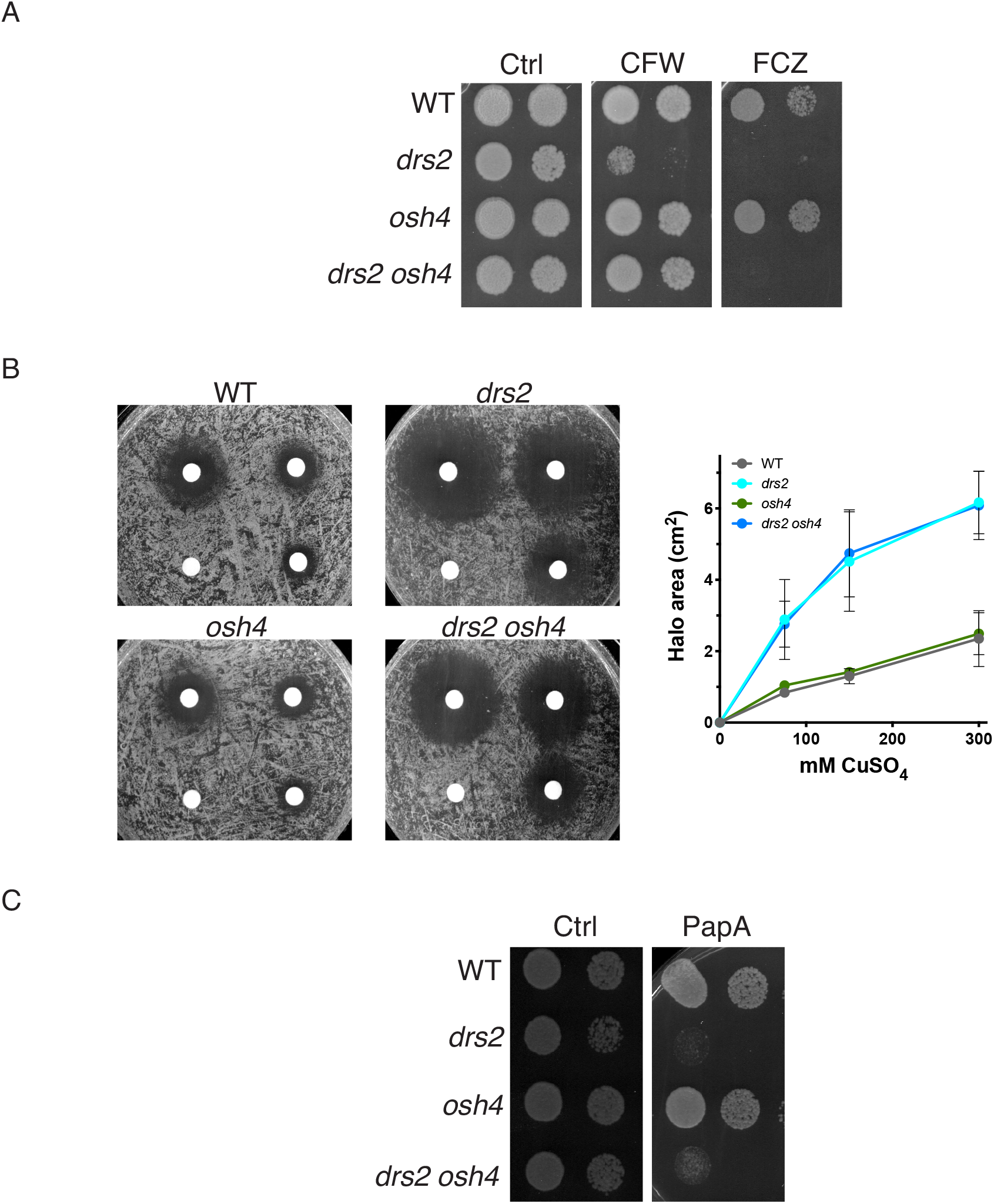
Deletion of *OSH4* does not rescue growth on fluconazole or copper in the *drs2* mutant. A) Deletion of *OSH4* rescues growth on calcofluor white but not that on fluconazole in the *drs2* mutant. Serial dilutions of indicated strains as in Fig. 2B were spotted on YEPD media (Ctrl) containing 25 μg/ml calcofluor white (CFW) or 5 μg/ml fluconazole (FCZ). Images were taken after 2 days. Similar results were observed in 2 experiments. B) Deletion of *OSH4* does not restore growth on copper in the *drs2* mutant. The indicated strains, WT (PY4861), *drs2/drs2* (*drs2*, PY3375), *drs2/drs2 osh4/osh4* (*drs2 osh4*, PY5539) and *osh4/osh4* (*osh4*, PY3974), were spread on YEPD and filter disks contained 10 μl of 0, 75, 150, or 300 mM CuSO_4,_ were added as in [35]. The zone of growth inhibition surrounding the filter discs was recorded after 1 day at 30°C and graphs represent averages of 3 independent experiments. C) Deletion of *OSH4* does not restore growth on papuamide A of the *drs2* mutant. Serial dilutions of indicated strains, WT (PY4861), *drs2/drs2* (*drs2*, PY3375), *drs2/drs2 osh4/osh4* (*drs2 osh4*, PY5539) and *osh4/osh4* (*osh4*, PY3974), were spotted on YEPD media (Ctrl) containing 1 μg/ml papuamide A (PapA) and images were taken after 2 days. Similar results were observed in 2 experiments.

As shown above, using the LactC2 reporter, we observed that the distribution of PS was altered during hyphal growth in the *drs2* mutant [36]. Papuamide A (PapA) is a depsipeptide toxin that binds PS in the outer leaflet of the PM [66]. In *C. albicans,* the *cho1* deletion mutant, which lacks the PS synthase, is less sensitive to PapA than WT cells [67]. Figure 9C shows that the *drs2* mutant was hypersensitive to PapA, compared to the wild-type strain, reflecting increased PS in the PM outer leaflet. Interestingly, the *drs2 osh4* mutant did not grow on PapA, similar to *drs2*, while the *osh4* mutant grew similar to the wild-type (Figure 9B). These results indicate that in this *drs2 osh4* mutant, the PM PS bilayer asymmetry is not re-established.

### The distinct roles of Drs2 can be associated with different localizations

Our data indicate that while filamentous growth was restored in the *drs2* mutant by *OSH4* deletion, growth on fluconazole and papuamide A was not. On the other hand, Figure 4 shows that, in *C. albicans*, Drs2 localizes to different structures, including the SPK. Hence, we sought to determine whether a specific localization of Drs2 is critical for distinct functions. To address this question, we used a synthetic physical interaction (SPI) approach to stabilize Drs2 at the SPK [50]. To generate a mutant with Drs2 restricted/stabilized at the SPK, we used a strain expressing a GFP nanobody (GNB) fused to one copy of Mlc1-iRFP, together with Drs2GFP as the Drs2 sole copy. Figure 10A (bottom panel) shows that, in such a mutant, Drs2 and Mlc1 co-localized at the SPK, with the Drs2 signal strikingly increased in the strain expressing Mlc1-GNB, compared to a control strain (top panel), consistent with Drs2 stabilized at this structure. In this strain expressing Mlc1-GNB, Drs2 was not detected along the PM, compared to what we observed in the absence of the GNB construct (Figure 4 and Figure 10A, top panel). In such a mutant expressing Mlc1-GNB, invasive filamentous growth was not altered (Figure 10B), with 85% of the cells forming hyphae in serum-containing liquid media after 90 min. In contrast, this mutant did not grow on fluconazole or papuamide A (Figure 10C). These data indicate that different roles of Drs2 are associated with distinct localizations, and suggest that *C. albicans* sensitivity to papuamide A is associated with Drs2 localization at the PM.

**Figure 10:**
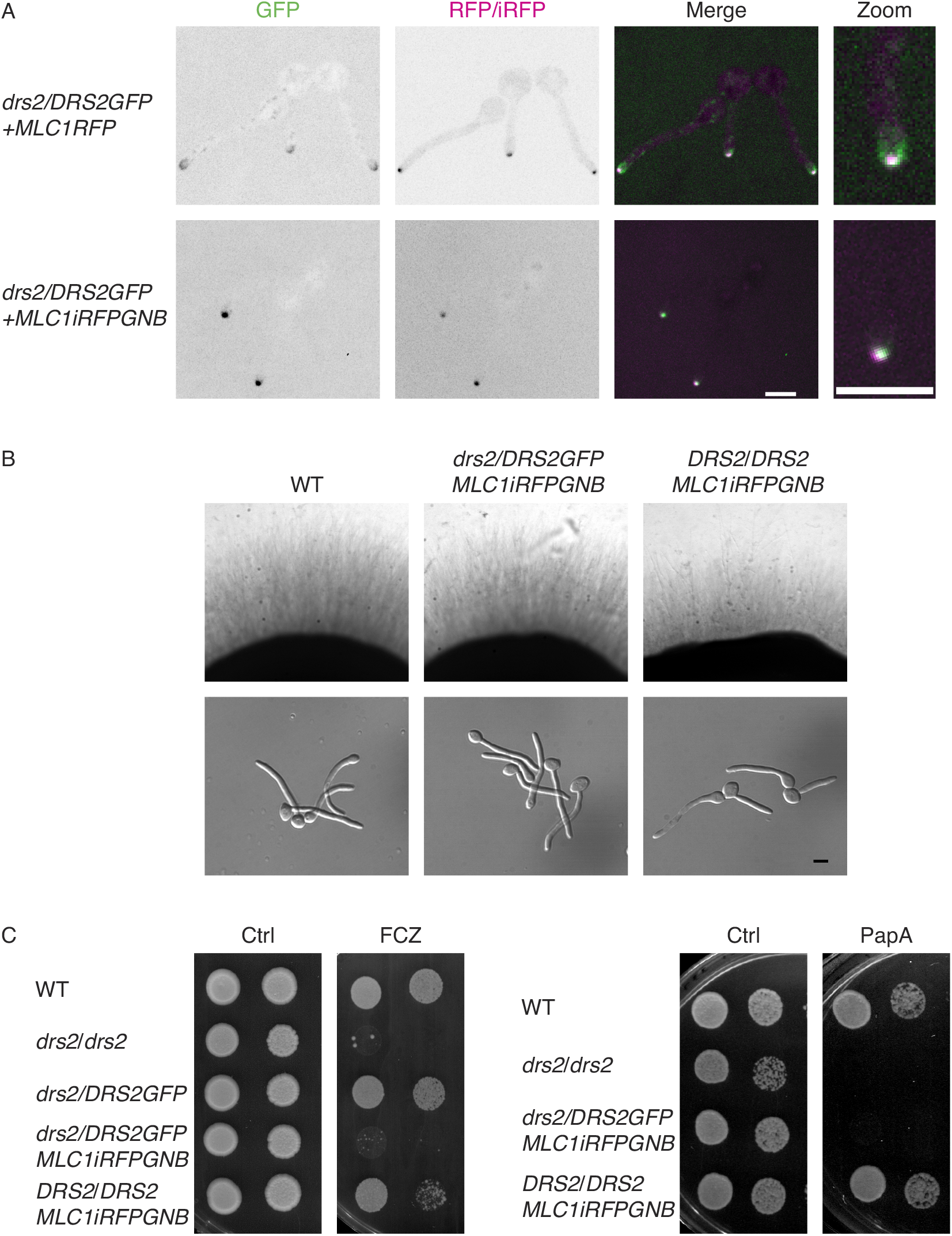
Stabilization of Drs2 at the Spitzenkörper alters growth on fluconazole and papuamide A. A). Synthetic physical interaction (SPI) between Drs2 and Mlc1 stabilizes Drs2 at the Spitzenkörper. Representative images (central z-sections) of cells expressing Drs2-GFP, as a sole copy of Drs2, with either Mlc1RFP (PY4928) or Mlc1-iRFP-GNB (PY5885) are shown; iRFP stands for near-*i*nfra*r*ed *f*luorescent *p*rotein and GNB for GFP nanobody. Right panels are enlargement of merge images. B) Stabilization of Drs2 at the SPK does not alter invasive filamentous growth. The indicated strains, WT (PY4861), *drs2/DRS2GFP MLC1-iRFP-GNB* (PY5885) and *DRS2/DRS2 MLC1-iRFP-GNB* (PY5385) were grown on agar-containing serum media and images were taken after 6 days (top panels) or grown in the presence of serum for 90 min (bottom panels). The percent of filamentous cells was 85-86% for the wild-type, *drs2/DRSGFP MLC1-iRFP-GNB* and *DRS2/DRS2 MLC1-iRFP-GNB* strains; 2 experiments, *n* ∼ 100 cells. C) Stabilization of Drs2 at the SPK alters growth on fluconazole and papuamide A. Serial dilutions of WT (PY4861), *drs2/drs2* (PY3375), *drs2/DRS2GFP* (PY4665), *drs2/DRS2GFP MLC1-iRFP-GNB* (PY5885) and *DRS2/DRS2 MLC1-iRFP-GNB* (PY5385) strains spotted on YEPD media (Ctrl) containing either 2.5 μg/ml fluconazole (FCZ) or 2 μg/ml papuamide A (PapA) and images were taken after 2-3 days.

Given that deletion of *OSH4* restored filamentous growth in a *drs2* mutant, but not growth on fluconazole or papuamide A, we investigated the effect of recruiting Osh4 to the SPK in these growth conditions. In *S. cerevisiae*, it was shown that the majority of Osh4 was cytoplasmic, with some punctae concentrated in small buds and mother–daughter necks [68]. In *C. albicans* hyphal cells, we did not detect a distinct localization of Osh4 (Figure 11A, top left panel). However, in cells expressing both Mlc1-GNB and Osh4-GFP, as the sole Osh4 copy, Osh4 co-localized with Mlc1 at the SPK, indicating that Mlc1-GNB recruited this LTP (Figure 11A). Such a recruitment of Osh4 in a wild-type strain background did not alter filamentous growth, yet it recovered filamentous growth in liquid media and partially recovered that on solid media in a *drs2* strain background (Figure 11B). In contrast, growth on papuamide A was not restored (Figure 11C). Together, these data indicate that recruitment of Osh4 to the SPK has a similar affect as deleting *OSH4*, in *drs2* cells for filamentous growth.

**Figure 11:**
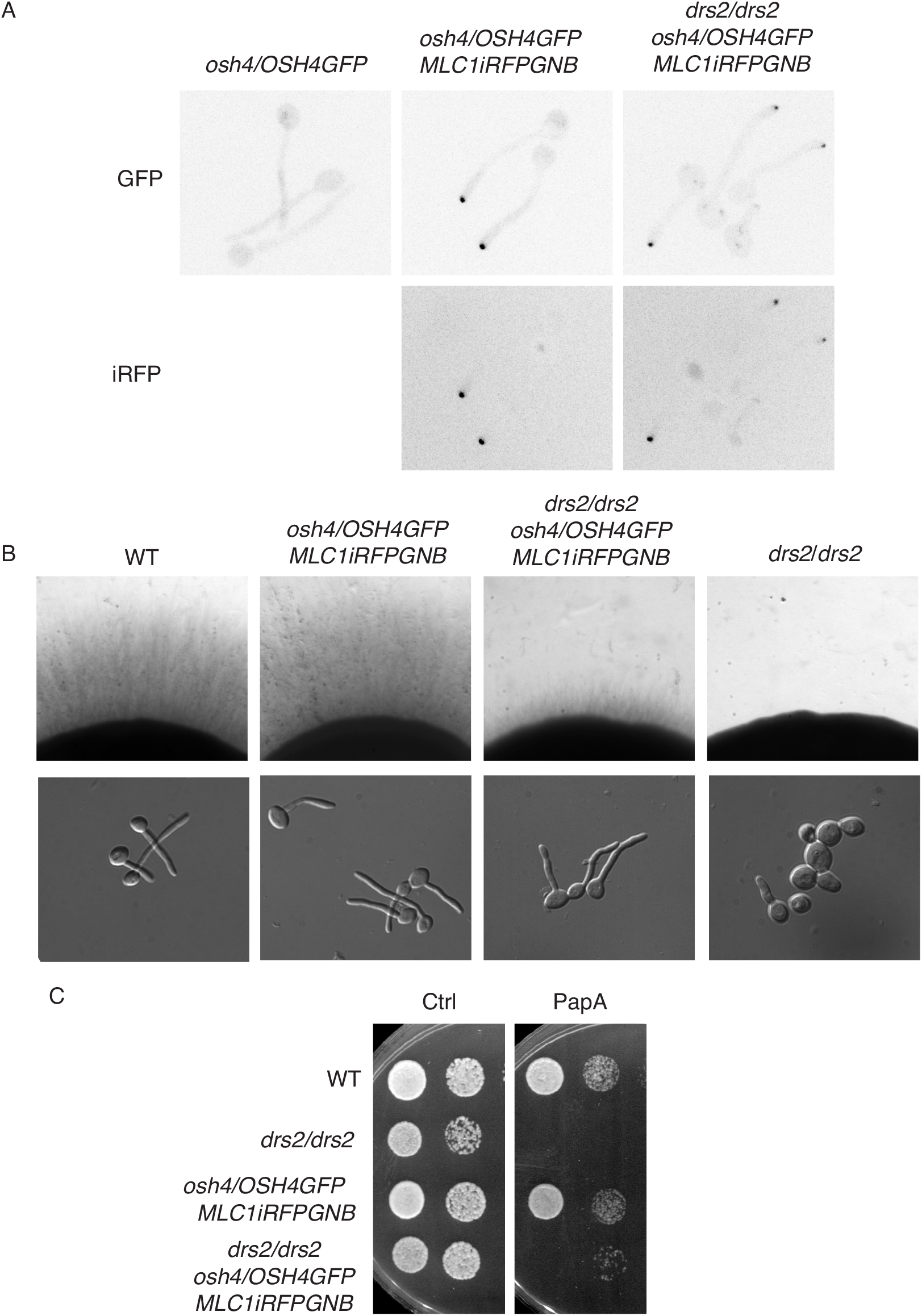
The recruitment of Osh4 to the Spitzenkörper in *drs2* phenocopies *drs2 osh4*. A). Synthetic physical interaction between Osh4 and Mlc1 recruits Osh4 to the Spitzenkörper. Representative images (Max projections) of cells expressing Osh4-GFP, as a sole copy of Osh4, without (PY5875) or with Mlc1-iRFP-GNB, in the wild-type (PY5931) or the *drs2* deletion (PY6447) strains are shown. B) Recruitment of Osh4 at the SPK restores invasive filamentous growth in a *drs2* deletion strain. Indicated strains, wild-type (PY4861), *drs2/drs2* (PY3375), *drs2/drs2 osh4/OSH4GFP MLC1-iRFP-GNB* (PY6447) and *DRS2/DRS2 osh4/OSH4GFP MLC1-iRFP-GNB* (PY5931) were grown on agar-containing YEPD with serum and images were taken after 6 days (top panels) or grown in the presence of serum for 90 min (bottom panels). At 90 min, the percent of filamentous cells was on average 85%, 78%, 78% and 22% for PY4861, PY5931, PY6447 and PY3375 strains, respectively; 2-3 independent experiments with *n* ∼ 100 cells. C) Recruitment of Osh4 at the SPK does not restore growth on papuamide A in the *drs2* mutant. Serial dilutions of indicated strains as in Fig. 11B, were spotted on YEPD media (Ctrl) containing 2 μg/ml papuamide A (PapA) and images were taken after 2-3 days.

## Discussion

Our results show that Drs2 is critical for *C. albicans* invasive filamentous growth and that a *drs2* deletion mutant exhibits increased sensitivity to calcofluor white and fluconazole, as well as papuamide A. The *drs2* mutant also has an altered PS, PI(4)P and ergosterol distribution, but not that of PI(4,5)P_2_. The requirement for Drs2 in invasive filamentous growth, as well as growth in the presence of the chitin binding dye calcofluor white, can be specifically bypassed by the deletion of the lipid transfer protein Osh4, but not by that of other Osh proteins. Deletion of *OSH4* in *drs2* also recovers the PI(4)P distribution and significantly the PS distribution, but not growth on papuamide A, indicating that the level of PS in the PM outer leaflet remains altered. In *C. albicans*, Drs2 localizes to different compartments, *i.e.* the Golgi, the Spitzenkörper and the PM. We propose that distinct roles of Drs2 are associated with different localizations.

### Role of Drs2 in plasma membrane organization

In *S. cerevisiae*, it was shown that the *DRS2/DNF1-3* genes have some redundancy during budding growth [20]. Similarly, in the filamentous fungus *A. nidulans*, a double deletion of the Dnf1 and Drs2 homologs (DnfA and DnfB, respectively) is lethal [25, 69], indicating that these flippases are redundant, in particular for growth. Yet, they appear to also have unique functions and/or localizations in different fungi. For example, in contrast to the case in *C. albicans*, deletion of the Drs2 homolog DnfB alone does not drastically alter *A. nidulans* hyphal growth [25]. On the other hand, while ScDrs2 remains associated to the Golgi apparatus, even during mating in which Dnf1-3 localize to the shmoo tip [24], AnDnfB localizes to the SPK [25]. We show here that CaDrs2 localizes to the filament apex (SPK and PM in addition to the Golgi apparatus, and our data indicate that these distinct localizations are associated with different Drs2 functions. Indeed, upon stabilization of Drs2 at the SPK, in which this flippase is no longer detected at the PM, cells are similar to that of the wild-type for invasive filamentous growth and calcofluor white sensitivity, yet unable to grow on papuamide A. Furthermore, deletion of Osh4, which restores hyphal growth and calcofluor white sensitivity, does not restore *drs2* growth on papuamide A, hence the level of PS in the PM external leaflet. Copper binds PS with high affinity and a *drs2* mutant is hypersensitive to copper, linking PS and copper sensitivity [35]. Given that *drs2 osh4* grows similar to *drs2* on copper, we suggest that the PM localization of Drs2 is required for membrane organization, *via* PS flippase activity, ultimately regulating PS asymmetry.

### Role of lipids in invasive filamentous growth

A *drs2* deletion mutant is defective in invasive filamentous growth in response to serum, and deletion of *OSH4* can specifically overcome these defects. Upon serum exposure, the *drs2* cells exhibit an alteration in the PM lipid distribution, *i.e.* PS, PI(4)P and ergosterol, which is at least partially compensated by *OSH4* deletion, raising the question of the respective contribution of these lipids to hyphal growth.

Sterol-rich membrane domains, have been visualized, by using filipin, at the tips of mating projections in *S. cerevisiae* [70], at cell poles and division site in *S. pombe* [71], as well as at the hyphal tips of *C. albicans* [58] and *A. nidulans* [72], where they are thought to contribute to polarized growth. In *S. cerevisiae drs2* cells, 20% of ergosterol appears to mislocalize to internal membranes, as assessed by filipin staining and DHE fluorescence, and deletion of *KES1* suppresses the accumulation of intracellular ergosterol in this *drs2* mutant [73]. Our data, using the D4H reporter for free ergosterol within the cell, indicates that 96 ± 7% and 78 ± 12% of cells had multiple internal punctae in the *drs2* and *drs2 osh4* mutants, respectively, compared to 1 ± 2% and 9 ± 6% in wild-type and *osh4* cells, respectively, indicating that removal of Osh4 did not substantially suppress internal ergosterol accumulation in *C. albicans drs2* cells. Polarization of ergosterol to the growth tip also appears similar in the *drs2* and *drs2 osh4* mutants, at a level intermediate between the wild-type and *osh4* cells. Whether this polarized ergosterol signal is associated with the apical cortex and/or the SPK is unclear. In *S. pombe*, it was shown that ergosterol associated D4H internal signals, which are enriched upon Arp2/3 inhibition, correspond to membrane-enclosed compartments, referred to as sterol-rich compartments [60]. Both this ergosterol movement to internal structures and the anterograde transport appear to be independent of vesicular trafficking [60]. Although the mechanism of sterol transfer is still to be determined, it is unlikely that the filamentous growth defect in *drs2* directly results from the observed alteration of ergosterol distribution.

In *C. albicans*, the phosphatidylinositol 4-kinase *stt4* deletion mutant, in which PM PI(4)P is no longer detectable, and the PS synthase *cho1* mutant, which has little to no PS, exhibit defective filamentous growth and increased sensitivity to the antifungal drug caspofungin [9, 74]. Both these mutants have increased exposure of cell wall β(1,3)-glucan and, given that 50% of PS is still detectable at the PM of *stt4*, we proposed that PM PI(4)P levels are associated with virulence by masking of cell wall β(1-3)-glucan [74]. In contrast to these mutants, the *drs2* mutant grows similar to the wild-type cells on caspofungin and congo red, making it unlikely that the glucan synthase pathway is affected. As PM PS and PI(4)P levels are not drastically altered compared to the *cho1* and *stt4* deletion mutants, how the virulence is altered in *drs2* is unclear. Similar to the *drs2* mutant, the ratio of mean PM PS to internal PS decreases in a *C. albicans stt4* mutant, with increased internal PS levels compared to wild-type cells [74], yet, as for *stt4*, our results are inconsistent with a correlation between PS distribution and filamentous growth defect in the *drs2* mutant. Indeed, while invasive filamentous growth was recovered upon deletion of Osh4, the PM to internal PS ratio was only partially recovered, and deletion of Osh4 did not recover growth on papuamide A. Rather, PM PI(4)P appears to be critical. In *S. cerevisiae*, the cold-sensitive growth of a *drs2* deletion mutant is partially suppressed by deletion of either the Osh4 homolog, Kes1, or the PI(4)P phosphatase Sac1 [31]. In *C. albicans*, deletion of Sac1 results in cells with altered PM PI(4)P gradient, as well as filamentous growth [56, 64]. While the apical PI(4)P gradient as well as filamentous growth are restored upon deletion of Osh4 in *drs2* cells, deletion of Sac1 substantially affects growth in *drs2,* presumably by increasing the level of PM PI(4)P. Together, these results strongly suggest that the PM PI(4)P gradient, rather than the level of PI(4)P, is critical for filamentous growth.

### Role of Drs2 in cell wall integrity and fluconazole sensitivity

Lipid transport *via* vesicular and non-vesicular trafficking is well established [75] and PS flipping activity of Drs2 is critical for the vesicular transport of a number of PM proteins. For example, ScDrs2 is required at the Golgi for efficient segregation of cargo into exocytic vesicles, as the PM proteins Pma1 and Can1 are missorted to the vacuole in *drs2Δ* cells, and deletion of *KES1* suppresses this defect [73]. Hence, it is likely that the hypersensitivity of the *drs2* mutant both for calcofluor white and fluconazole results from an alteration of membrane traffic in specific pathways. Interestingly, the *drs2* mutant exhibits increased sensitivity to calcofluor white but not to caspofungin, strongly suggesting that the chitin synthase pathway is selectively altered. Consistently, it was proposed that PS translocation to cytosolic leaflet of the Golgi by a flippase is critical for the *S. cerevisiae* chitin synthase (Chs3) trafficking [76]. The growth defect of *drs2* on fluconazole may result from a mislocalization of multi-drug transporters [39] and, consistent with such a scenario, we observe that ∼ 80% of the *drs2* cells had internal Cdr1 signal. Given that Drs2 is critical for fluconazole sensitivity, it would be interesting to investigate the importance of such a flippase in pathogenic non *albicans* Candida species, known for their resistance to the antifungal drug, such as *C. glabrata* or *C. auris* [77].

In summary, altogether our results are consistent with the notion that Drs2 down-regulates Osh4 activity [73], as wild-type filamentous growth and cell wall integrity, were recovered in *drs2* either by deleting or targeting Osh4 to the SPK. However, the converse, *i.e.* Osh4 down-regulating Drs2 activity, appears less critical for *C. albicans* filamentous growth. Furthermore, our data suggest that Osh4 does not function together with Drs2 at the SPK for filamentous growth, as targeting Osh4 to this organelle has the same effect as deleting Osh4. Strikingly, our data show that the genetic interaction between two different lipid transporters, identified in *S. cerevisiae* budding growth [31], is also observed in a distinct growth state in *C. albicans*, which diverged from *S. cerevisiae* **∼**250 million years ago [78]. Whether this is a conserved feature of the fungal kingdom is an attractive possibility, and filamentous fungi, with large diversity in size and growth rate, represent useful models to understand how membrane expansion is regulated by lipid transport mechanisms.

## Materials and Methods

### Growth conditions

Yeast extract-peptone dextrose (YEPD) or synthetic complete (SC) medium was used and strains were grown at 30°C, unless indicated otherwise. Filamentous growth induction was carried out as described previously either with 50% serum or 75% serum [79]. For filipin staining experiments, 10% of serum was used. Growth on YEPD plates containing Congo red, calcofluor white, caspofungin or fluconazole was examined as described [80]. Copper sensitivity was investigated as described [30]. Congo red, calcofluor white, filipin, hygromycin and fluconazole were from Fluka, Sigma-Aldrich, Saint Quentin Fallavier, France. Papuamide A was from University British Columbia.

### Strains and plasmids

Strains and oligonucleotides used are listed in Tables S1 and S2, respectively. All strains were derived from BWP17 [81]. The deletion mutants were generated by homologous recombination. Each copy was replaced by either *HIS1*, *URA3*, *ARG4*, *SAT1* or *HYGB*, using knockout cassettes generated by amplification of pGem-*HIS1*, pGem-*URA3*, pGem-*CdARG4*, pFa-*ARG4*, pFa-*SAT1* and pBH1S [81–84] with primer pairs *DNF1.P1/DNF1.P2, DNF2.P1/DNF2.P2, DNF2.P3/DNF2.P4 DNF3.P1/DNF3.P2, OSH4.P1/OSH4.P2*, *OSH7A.P1/OSH7A.P2*, *OSH7α.P1/OSH7α.P2, OSH2.P1/OSH2.P2, OSH3.P1/OSH3.P2* and *SAC1.P1/SAC1.P2*. The *drs2Δ/drs2Δ osh4Δ/osh4Δ, drs2Δ/drs2Δ osh2Δ/osh2Δ*, *drs2Δ/drs2Δ osh3Δ/osh3Δ*, *drs2Δ/drs2Δ osh7AΔ/osh7αΔ* and *drs2Δ/drs2Δ sac1Δ/sac1Δ* strains were generated from the *drs2Δ/drs2Δ* strain (PY3310 [36]) and the *dnf1Δ/dnf1Δ dnf2Δ/dnf2Δ* strain from the *dnf2Δ/dnf2Δ* strain (PY5804).

Plasmid *pDUP3*-*pADH1DNF2* was constructed by amplification from gDNA of the *DNF2* ORF, using primers with a unique AscI site at the 5’ end (DNF2.P5) and a unique MluI site at the 3’ end (DNF2.P6) and *pDUP3*-*pDRS2DRS2* by amplification from *pExp-pDRS2DRS2* [36] of the *DRS2* ORF together with 1 kb upstream and downstream, using primers with a unique SpeI site at the 5’ end (*DRS2.P1*) and a unique NotI site at the 3’ end (*DRS2.P2*). The fragments were subsequently cloned into *pDUP3-SAT1* [85], yielding to *pDUP3-pADH1DNF2* and *pDUP3-pDRS2DRS2,* respectively. To visualize Drs2, Dnf2 and Osh4, GFPγ was inserted by homologous recombination at the 3’ end of *DRS2*, *DNF2* or *OSH4* ORF in strains heterozygous for these genes, after amplification of GFPγ from the plasmid *pFA-GFPγ-HIS1* [86], using the primers *DRS2.P3/DRS2.P4, DNF2.P7/DNF2.P8* or *OSH4.P3/OSH4.P4*. *pExp-pACT1CRIBGFP* [48], was used to transform the WT (BWP17) and *drs2/drs2* (PY3310) strains. Cdr1-GFP, PH-FAPP1^[E50A,H54A]-^GFP, GFP-(PH^OSH2[H340R]^)_2_-GFP and GFP-PH_PLCδ1_-PH_PLCδ1_-GFP expressing strains were generated as described [56, 57]. pDUP5-mScarlet-CtRac1 [87] was used to transform the *drs2/DRS2GFP* (PY4665) and *dnf2/DNF2GFP* (PY5746) strains. Mlc1-mScarlet and Cdc10-mScarlet were generated by amplification of mScarlet-ARG4 from pFA-mScarlet-ARG4 [88], using the primer pairs *MLC1.P1/MLC1.P2* and *CDC10.P1/CDC10.P2*, respectively. mTurquoise-CtRac1 (mTrq-CtRac1) was constructed by replacing mScarlet in *pDUP5-pADH1-mScarlet-CtRac1* [87] with mTrq, and subsequently cloning mTrq-CtRac1 into *pDUP3* (Puerner, Bassilana & Arkowitz, in preparation). To visualize the distribution of phosphatidylserine, a fusion of GFP with the discoidin-like C2 domain of lactadherin (GFP-LactC2) was used, as previously described [36]. Strains expressing Mlc1-iRFP-GNB were generated by homologous recombination, using pFA-iRFP670-GNB-URA3, as in [50]. To visualize the distribution of ergosterol, we used filipin staining, as described [58], as well as the genetically encoded biosensor D4H [60]. D4H was amplified from plasmid pSM2244 [60], using primer pair D4H.P1 and D4H.P2 and cloned into plasmid *pExp-pACT1-mScarlet-CtRac1* [88], using AscI and MluI unique restriction sites, yielding *pExp-pACT1-mScarlet-D4H*. This plasmid was linearized with NcoI and integrated into the *RP10* locus. All other pExp plasmids were linearized with StuI and integrated into the *RP10* locus. pDUP3 and pDUP5 plasmids were digested with NgoMIV to release the cassette to be integrated into the *NEUTL5* locus.

Two independent clones of each strain were generated and confirmed by PCR. RT-PCR was also performed, where relevant, using the primers (GENE.pTm and GENE.mTm) listed in Table S2 or previously described [89] and RNA extraction was carried out using Master Pure yeast RNA extraction purification kit (Epicentre). All PCR amplified products were confirmed by sequencing (Eurofins MWG Operon, Ebersberg, Germany).

### Microscopy analyses

Colony and cell morphology imaging were performed as described [79]. Briefly, plates were incubated for 3–6 days prior to imaging with a Leica MZ6 binocular (x20) and cells (budding or serum-induced) were imaged by differential interference contrast with a microscope Leica DMR, using an ImagingSource DMK 23UX174 sCMOS camera.

Time lapses and fluorescent images were obtained using a spinning disk confocal microscope (inverted IX81 Olympus microscope with a 100X objective and a numerical aperture 1.45) and an EMCCD camera (Andor technology, UK). Z-stacks (images of 0.4 µm sections) acquired every 5 min, as described [56]. Maximum or sum intensity projections were generated from 21 z-sections with ImageJ software. Laser illuminations of 445 nm (Turquoise), 488 nm (GFP), 561 nm (mScarlet) and 670 nm (iRFP) were used. CRIBGFP distribution experiments were carried out as described [48]. For filipin analyses, widefield images were acquired on an inverted ZEISS Axio Observer Z1 microscope with a 100X (1.3 NA) objective. Huygens professional software version 18.04 (Scientific-Volume Imaging) was used to deconvolve z-stack images of cells expressing PH-FAPP1^[E50A,H54A]-^GFP. Bars are 5 µm.

Filament lengths and diameters were measured from DIC images, using image J software. Golgi cisternae were quantitated from deconvolved MAX projection images. Quantitation of PI4P distribution was performed on SUM projections, using the Matlab program HyphalPolarity [56]. Quantitation of PS PM and internal mean signals was performed on central z-sections, also using the Matlab program HyphalPolarity. Unless indicated otherwise, error bars represent the standard deviations. Statistical significance was determined with Student’s unpaired two-tailed *t* test.

### Virulence assays

HDC was induced in 10 Balb/C mice (Charles Rivers, Italy) per group by injecting the lateral tail vein with an inoculum of 5 x 10^5^ cells [90]. Animal body weight was monitored daily and animals were sacrificed by cervical dislocation when they had lost more than 20% of their weight.

### Ethics statement

Animal procedures were approved by the Bioethical Committee and Animal Welfare of the Instituto de Salud Carlos III (CBA2014_PA51) and of the Comunidad de Madrid (PROEX 330/14) and followed the current Spanish legislation (Real Decreto 53/2013) along with Directive 2010/63/EU.

**Table I:**
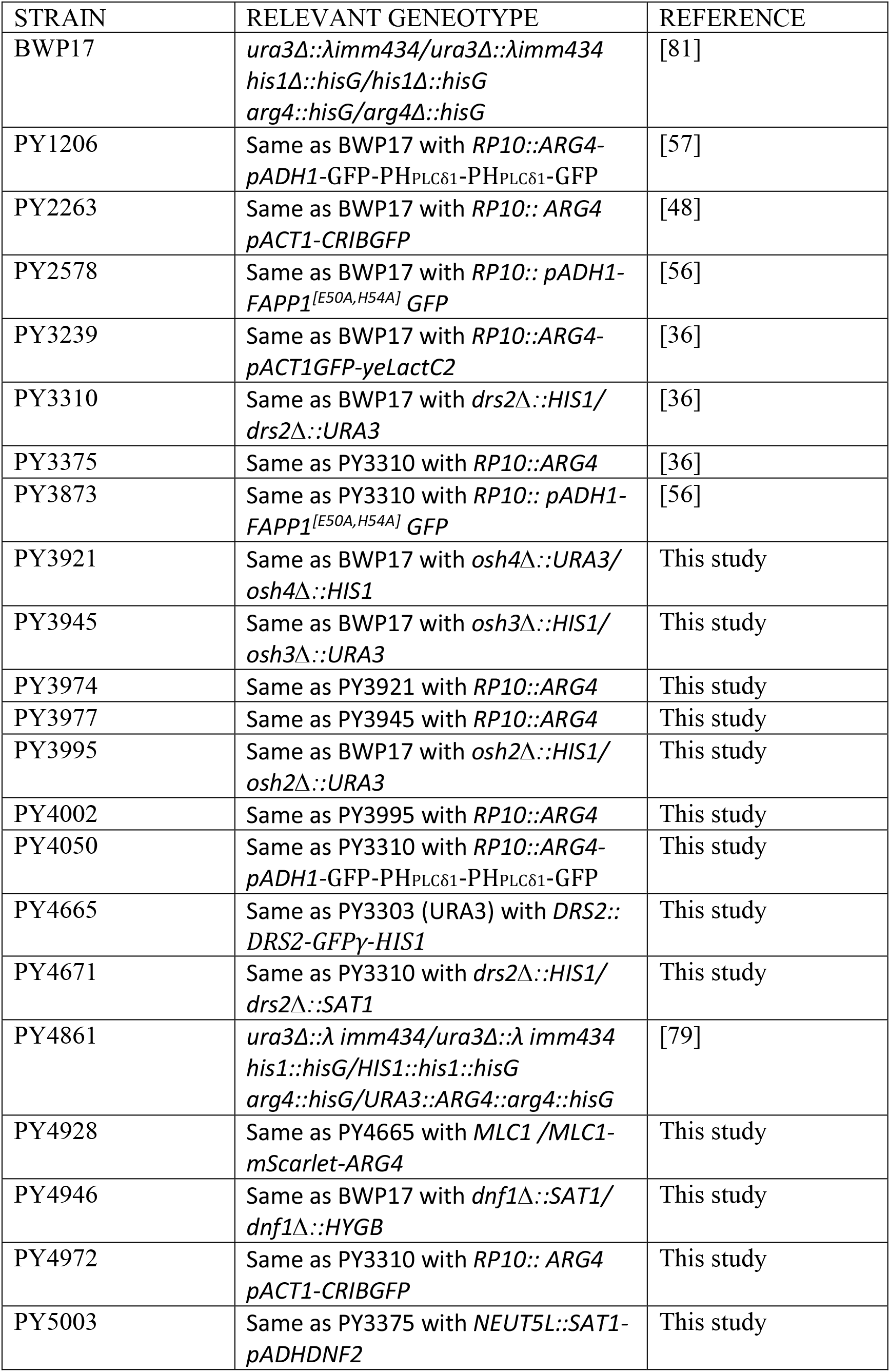
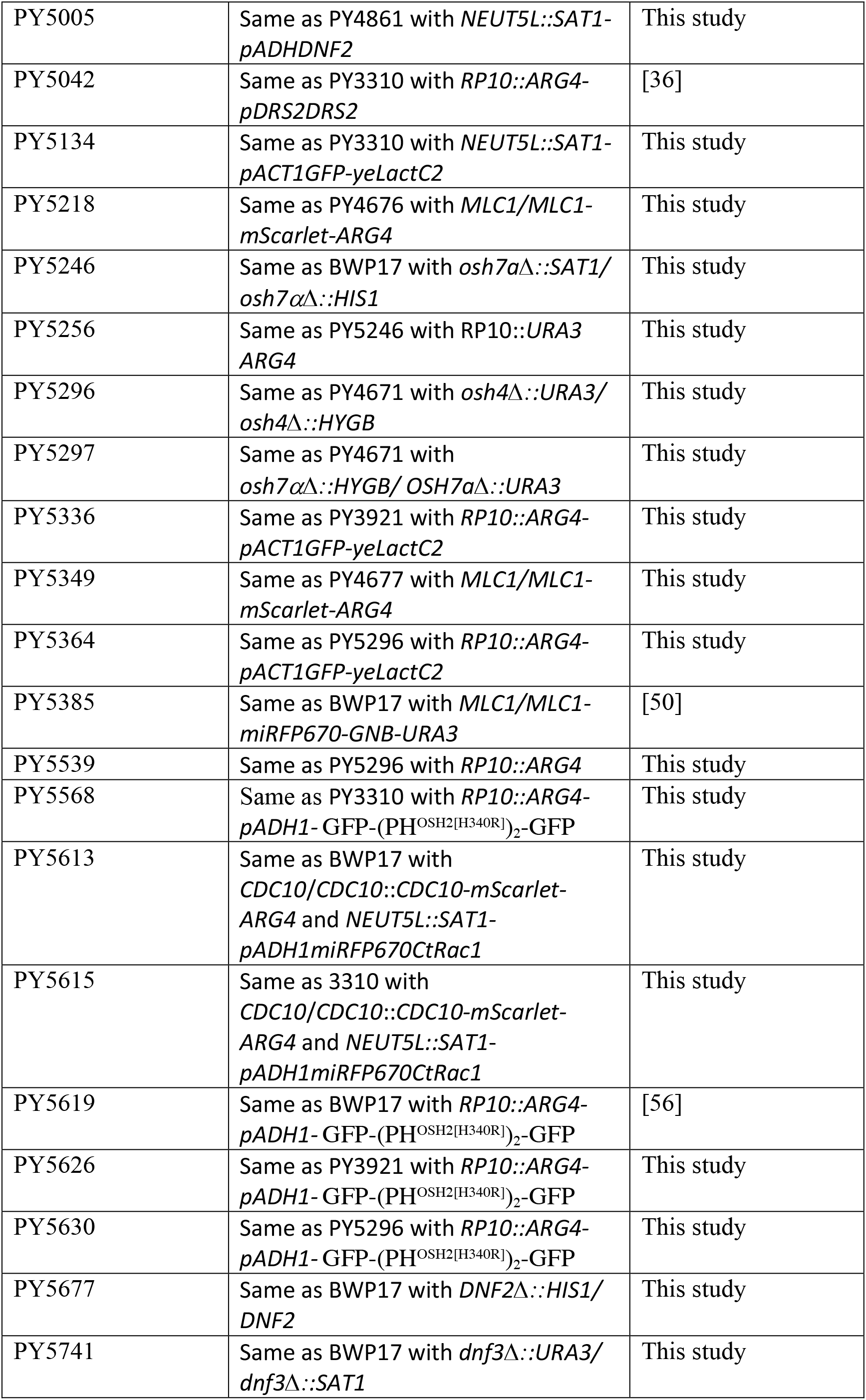
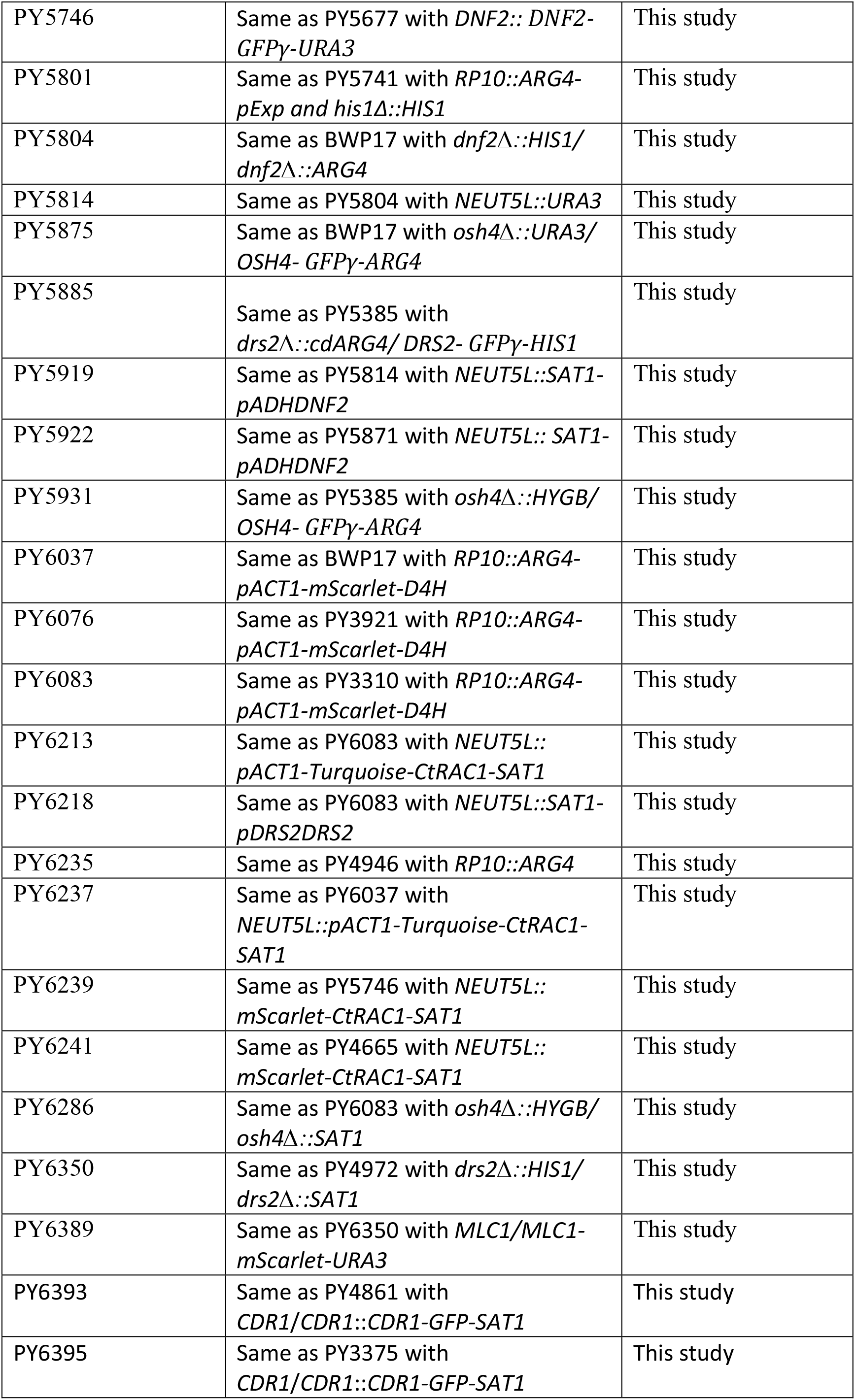
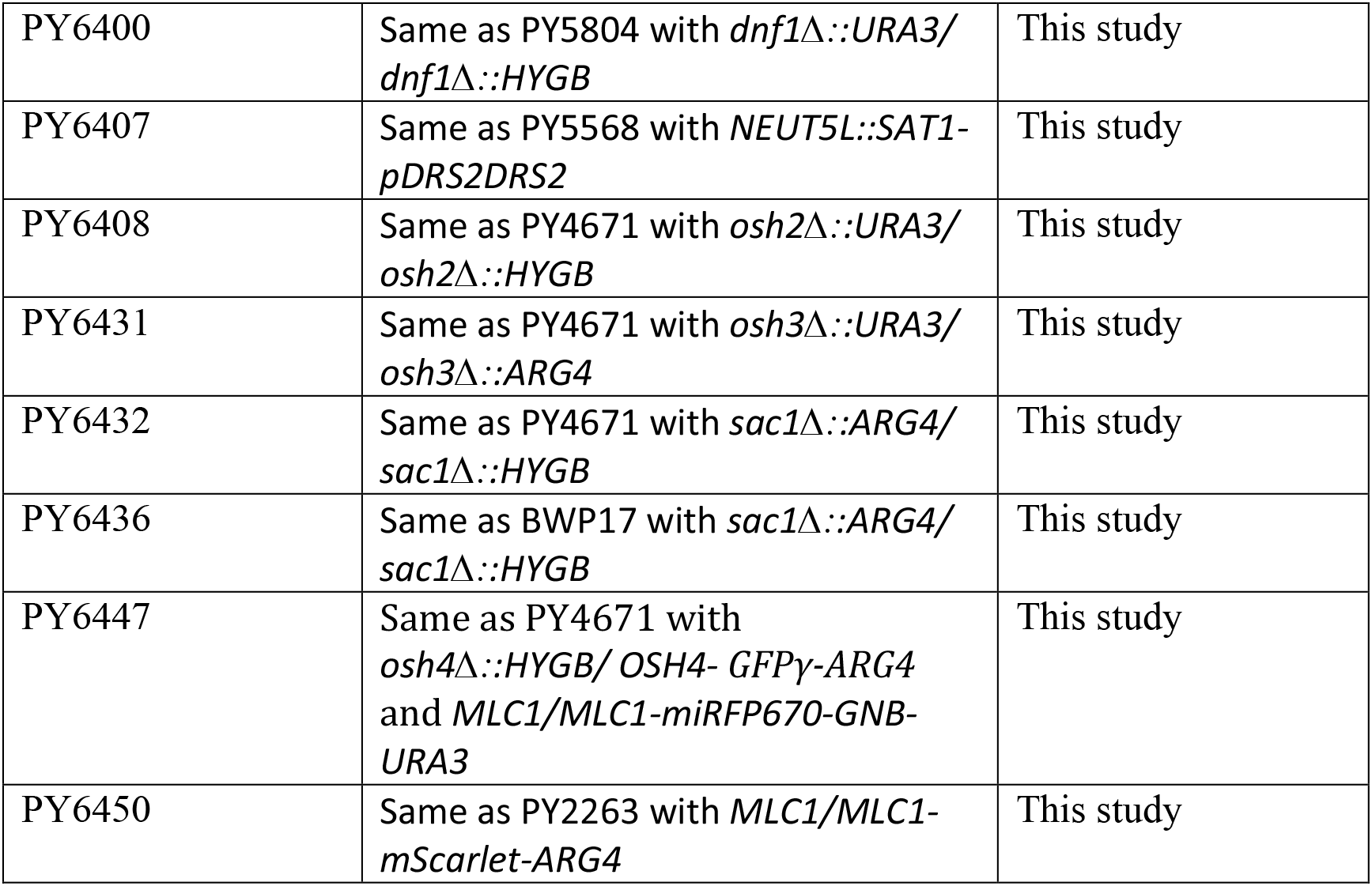
Strains used in the study.

**Table II:**
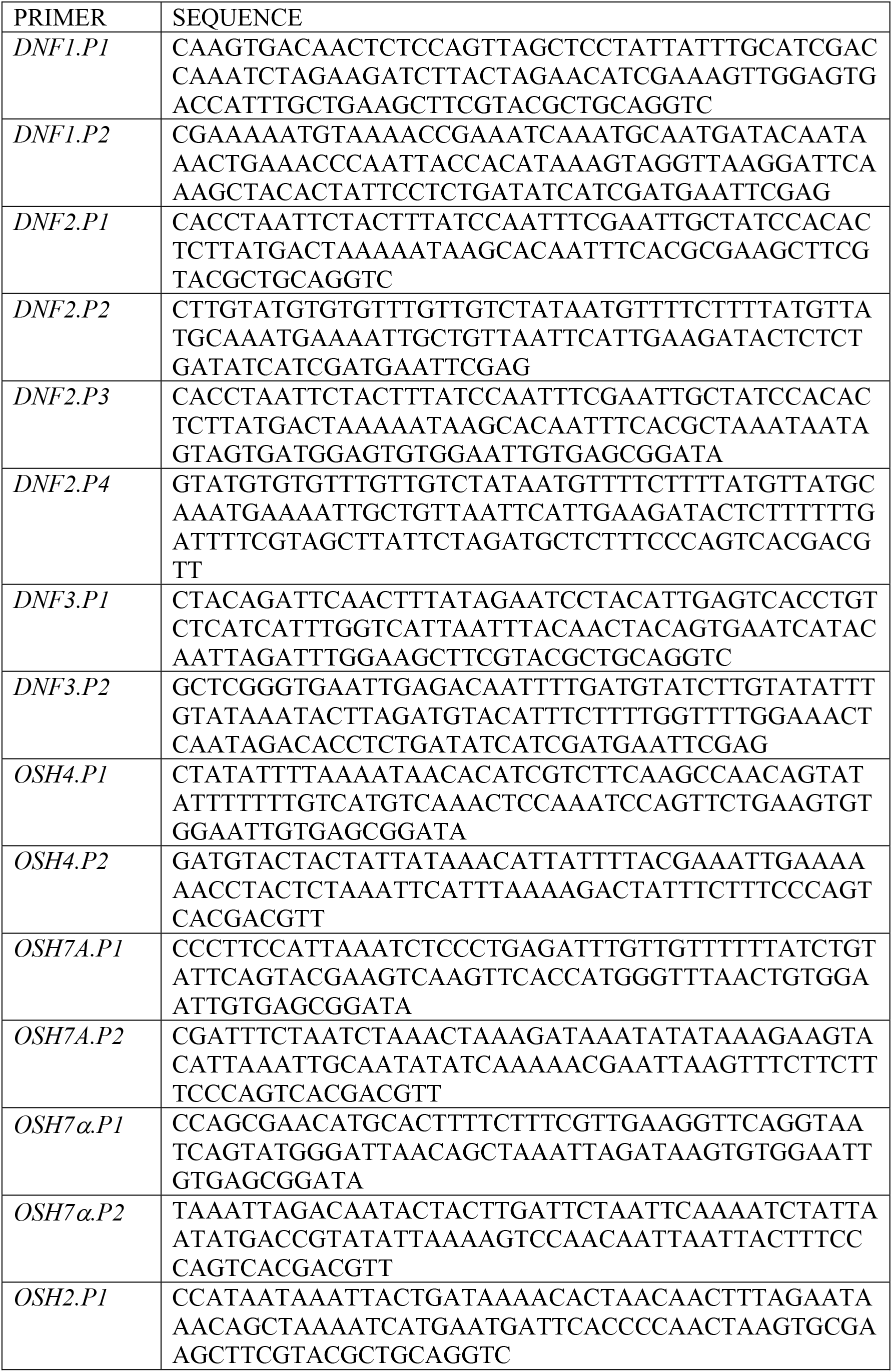
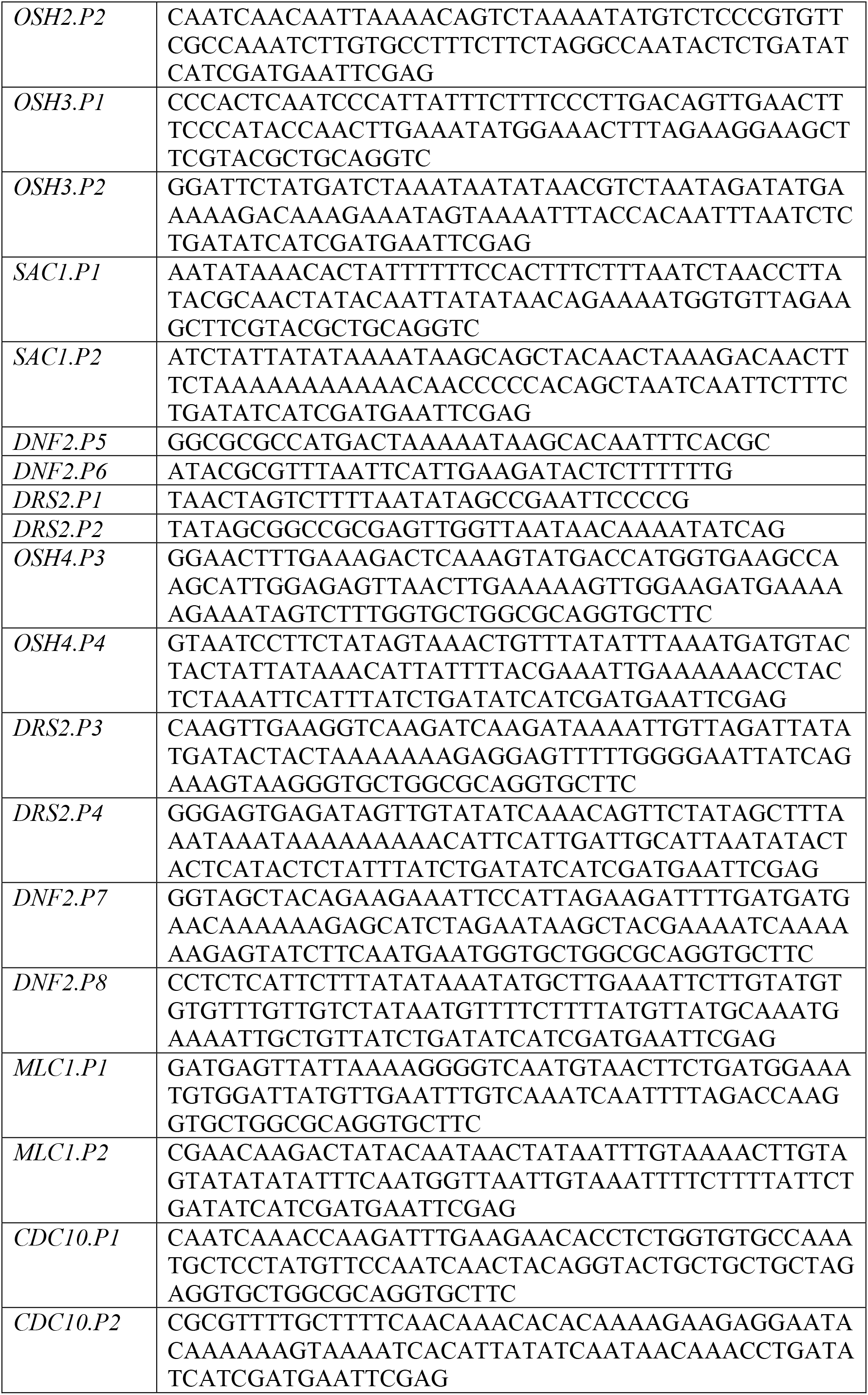
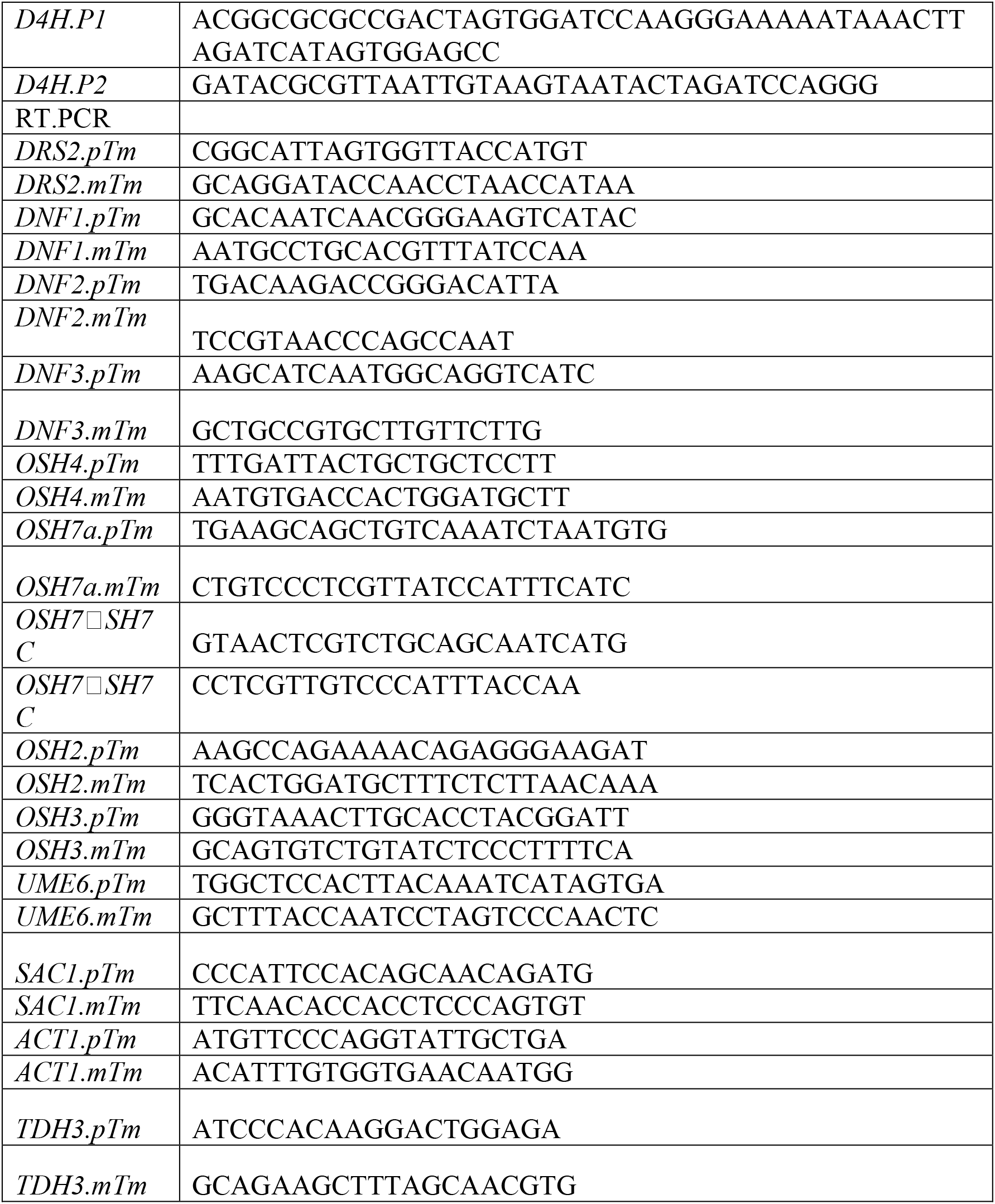
Primer sequences

## Acknowledgments

We thank J. Wendland, J.-C. Shieh and A. Mitchell for strains and plasmids. We thank the Platform Resources in Imaging and Scientific Microscopy facility (PRISM; M. Mondin, S. Lachambre, and B. Monterroso), and Microscopy Imaging Côte d’Azur (MICA) for microscopy support. This work was supported by the CNRS, INSERM, Université Côte d’Azur, and ANR (ANR-11-LABX-0028-01, ANR-16-CE13-0010-01 and ANR-19-CE13-0004-01) grants, and grant SAF2017-86192 from the Spanish Ministry for Science and Innovation.

