## Supplemental Figures S1-S3 for "Two distinct lipid transporters together regulate invasive filamentous growth in the human fungal pathogen *Candida albicans*"

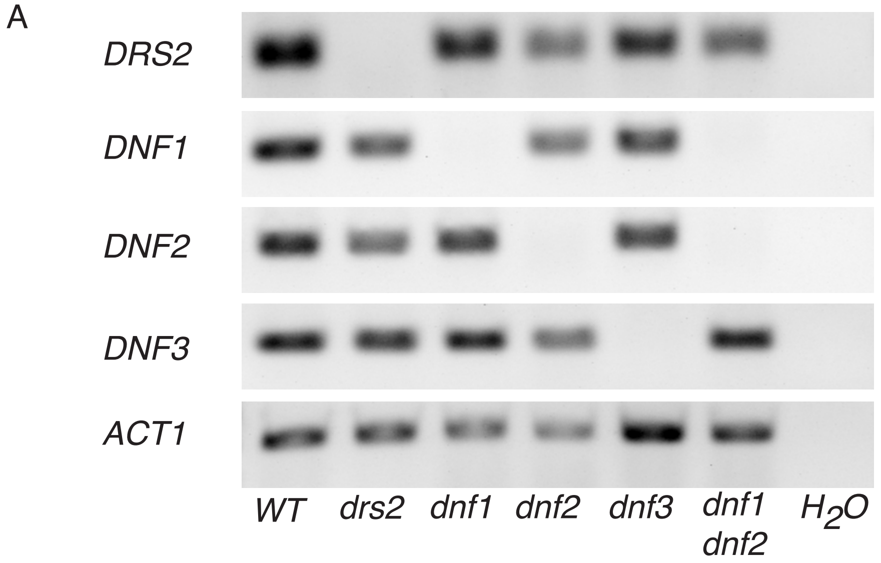


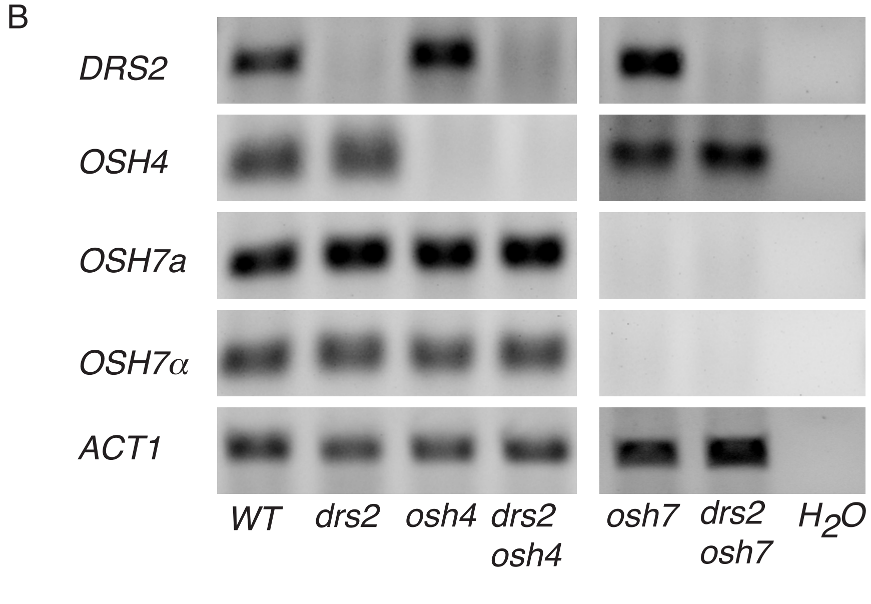


Figure S1: Flippase deletion mutant strains verification. A) *DRS2* and *DNF1-3* transcript levels. mRNA and cDNA were prepared from the indicated strains, as in Fig. 1A. *DRS2* and *DNF1-3* transcripts were determined by RT-PCR, using *DRS2*.pTm/*DRS2*.mTm (62 bp), *DNF1*.pTm/*DNF1*.mTm (73 bp), *DNF2*.pTm/*DNF2*.mTm (106 bp) and *DNF3*.pTm/*DNF3*.mTm (104 bp) primer pairs, respectively. Actin (*ACT1*) transcript levels (*ACT1*.pTm/*ACT1*.mTm primer pair) were used for normalization. B) *OSH4 and OSH7* transcript levels. mRNA and cDNA were prepared from the indicated strains, as in Fig. 7A. Transcripts were determined as in Figure S1A, using *OSH4*.pTm/*OSH4*.mTm (69 bp), *OSH7A*.pTm/*OSH7A*.mTm (84 bp), *OSH7*α.pTm/ *OSH7*α.mTm (65 bp) primer pairs, respectively.

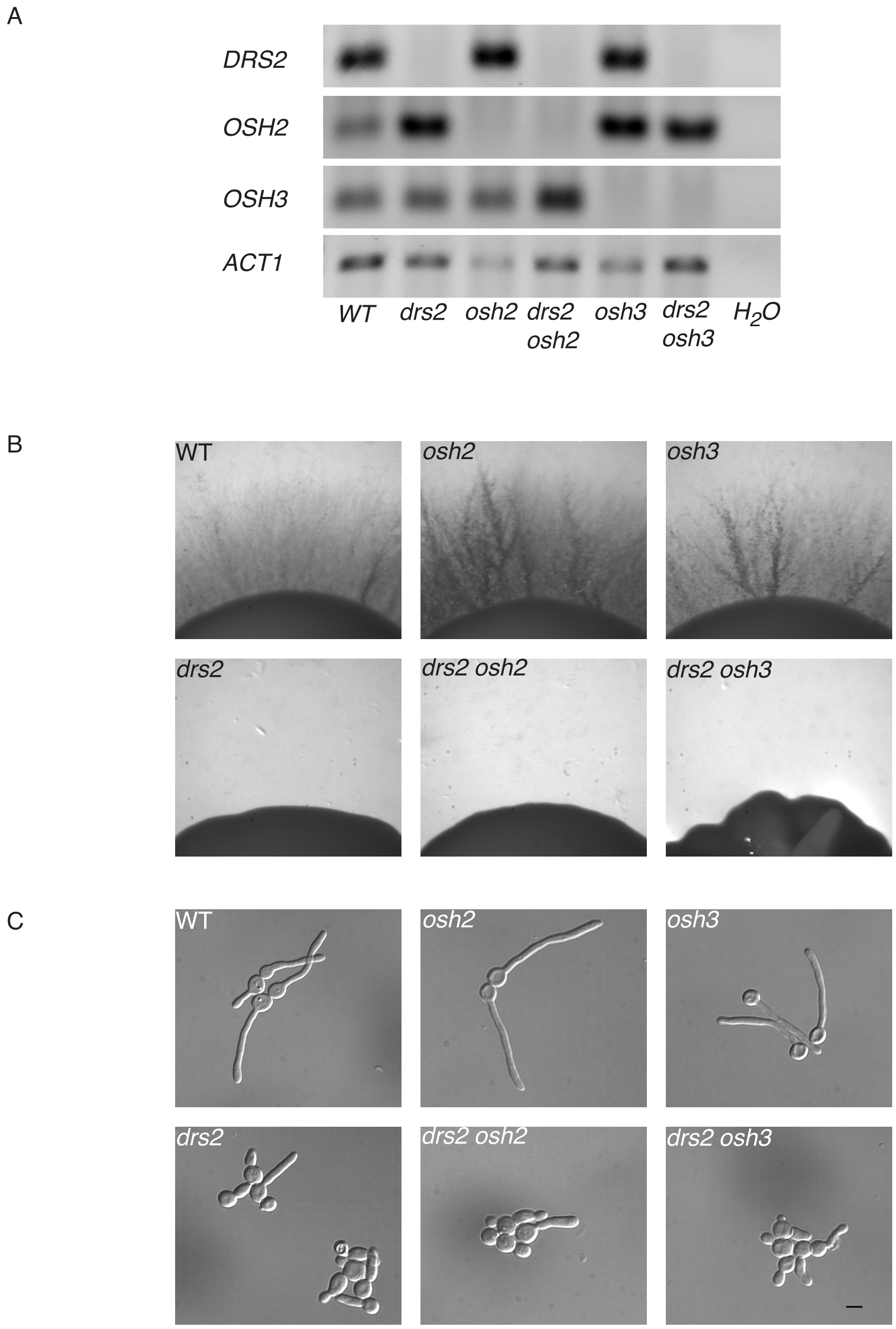


Figure S2: Deletion of *OSH2* and *OSH3* do not recover invasive filamentous growth in the *drs2* mutant. A) *DRS2* and *OSH2-3* transcript levels. mRNA and cDNA were prepared from the indicated strains, as in Fig. S3A. Transcripts were determined as in Fig. S1A, using *OSH2*.pTm/*OSH2*.mTm (74 bp), *OSH3*.pTm/*OSH3*.mTm (74 bp) primer pairs, respectively. B) Invasive growth is not restored in the *drs2* mutant upon deletion of *OSH2* or *OSH3*. The indicated strains, WT (PY4861), *osh2/osh2* (*osh2*, PY3977), *osh3/osh3* (*osh3*, PY4002), *drs2/drs2* (*drs2*, PY3375), *drs2/drs2 osh2/osh2* (*drs2 osh2,* PY6408) and *drs2/drs2 osh3/osh3* (*drs2 osh3*, PY6431), were grown on agar-containing serum media and images were taken after 6 days. Similar results were observed in 2 independent experiments. C) Hyphal growth is not restored in the *drs2* mutant upon deletion of *OSH2* or *OSH3*. At 90 min, the percent of filamentous cells, was determined from 3 independent biological samples for the WT, *drs2*, *osh2*, *osh3*, *drs2 osh2* and *drs2 osh3* strains; *n* ~ 100 cells.

A

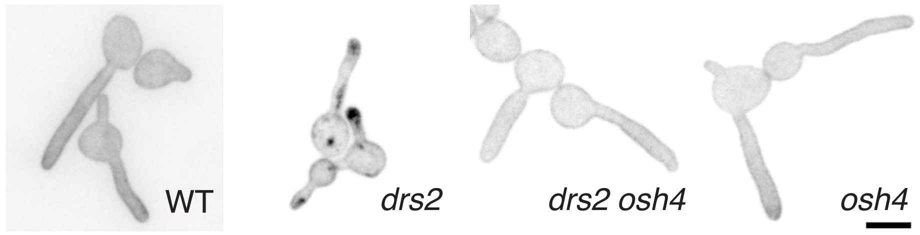


B

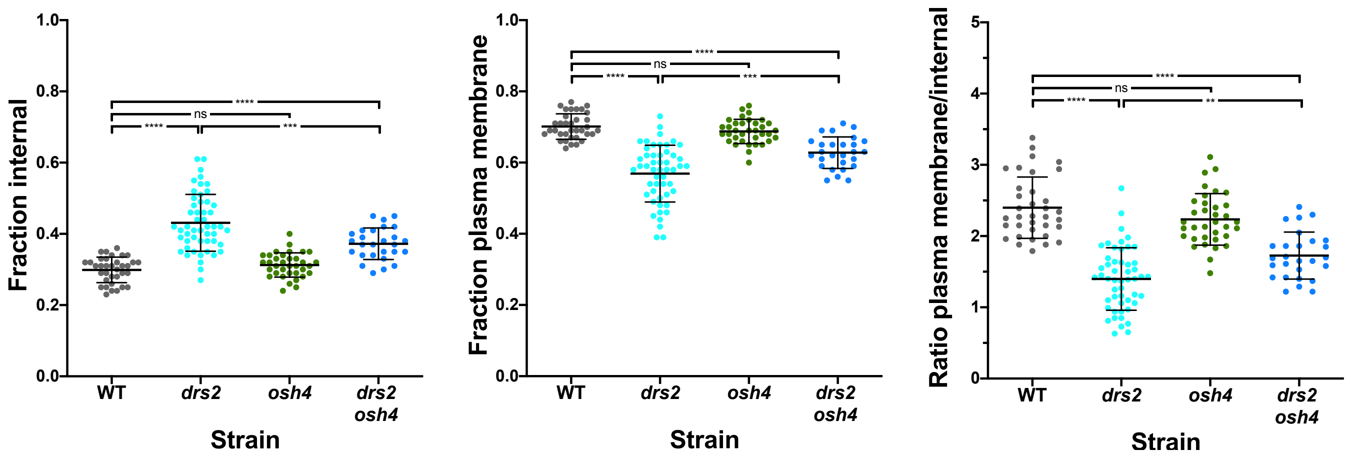


Figure S3: PS distribution is partially restored in the *drs2* mutant upon *OSH4* deletion. Indicated cells expressing GFP-LactC2, WT (PY3239), *drs2/drs2* (*drs2*, PY5134), *drs2/drs2 osh4/osh4* (*drs2 osh4*, PY5364) and *osh4/osh4* (*osh4*, PY5336) were incubated as in Fig. 5B, with sum projections of representative cells shown. The graphs represent the relative PM and internal PS fractions, as well as the ratio of the signal at the PM over the internal signal for the indicated strains ($n$ = 30-50 cells each), and bars mean ± the SD. ****, $P < 0.0001$; ***, $P < 0.001$; **, $P < 0.01$; ns, not significant.
